# Targeting Unique Ligand Binding Domain Structural Features Downregulates DKK1 in Y537S *ESR1* Mutant Breast Cancer Cells

**DOI:** 10.1101/2024.05.28.596307

**Authors:** K.S. Young, G.R. Hancock, E. Fink, A. Zigrossi, B. Flowers, D.A. Cooper, V.T. Nguyen, M. Martinez, K.S. Mon, M. Bosland, D. Zak, A. Runde, M.N. Sharifi, I. Kastrati, D.D.L. Minh, S. Kregel, S.W. Fanning

## Abstract

Resistance to endocrine therapies remains a major clinical hurdle in breast cancer. Mutations to estrogen receptor alpha (ERα) arise after continued therapeutic pressure. Next generation selective estrogen receptor modulators and degraders/downregulators (SERMs and SERDs) show clinical efficacy, but responses are often non-durable. A tyrosine to serine point mutation at position 537 in the ERα ligand binding domain (LBD) is among the most common and most pathogenic alteration in this setting. It enables endocrine therapy resistance by superceding intrinsic structural-energetic gatekeepers of ER hormone-dependence, it enhances metastatic burden by enabling neomorphic ER-dependent transcriptional programs, and it resists SERM and SERD inhibiton by reducing their binding affinities and abilities to antagonize transcriptional coregulator binding. However, a subset of SERMs and SERDs can achieve efficacy by adopting poses that force the mutation to engage in a new interaction that favors the therapeutic receptor antagonist conformation. We previously described a chemically unconventional SERM, T6I-29, that demonstrates significant anti-proliferative activities in Y537S ERα breast cancer cells. Here, we use a comprehensive suite of structural-biochemical, *in vitro*, and *in vivo* approaches to better T6I-29’s activities in breast cancer cells harboring Y537S ERα. RNA sequencing in cells treated with T6I-29 reveals a neomorphic downregulation of *DKK1*, a secreted glycoprotein known to play oncogenic roles in other cancers. Importantly, we find that DKK1 is significantly enriched in ER+ breast cancer plasma compared to healthy controls. This study shows how new SERMs and SERDs can identify new therapeutic pathways in endocrine-resistant ER+ breast cancers.

## INTRODUCTION

Over seventy percent of breast cancers are classified by their expression of the nuclear hormone receptor estrogen receptor alpha (ERα), encoded by the *ESR1* gene [1]. In these cases, the estrogenic steroid hormones bind to the receptor with high affinity, promoting transcriptional complex formation at enhancers and promoters that propels tumor cell proliferation, invasion, migration, and metastasis [2–5]. Hormone therapies target this transcription-driven pathology through direct and indirect effects on ERα [6, 7]. Aromatase inhibitors, such as anastrozole/arimidex, starve ERα of endogenous estrogens by preventing their conversion from androgens [8–10]. Direct ERα therapies, such as the selective estrogen receptor modulator (SERM) tamoxifen, achieve therapeutic endpoints by competitively binding to the hormone binding pocket within the ERα ligand binding domain (LBD) and favoring distinctive conformational ensembles that repopulate coregulator complexes to favor quiescent phenotypes [11, 12]. The second-line hormone therapy fulvestrant/faslodex, a selective estrogen receptor degrader/downregulator (SERD), competitively antagonizes transcription, but also induces proteasomal degradation by exposing buried hydrophobic LBD structural motifs to solvent [13, 14]. Although response to these primary targeted treatments in ER+ breast cancers is initially successful, over 30% of patients will relapse following 5 years of hormone therapy, highlighting the need to understand cellular mechanisms of therapy resistance [15].

*ESR1* missense mutations emerge after prolonged hormone therapy regiments and enable hormone therapy resistance by negating ERα’s hormone-dependence [16, 17]. Hotspot activating somatic missense mutations tyrosine 537 to serine (Y537S) and aspartic acid 538 to glycine (D538G) together account for >50% of identified mutants. Both mutations enable the formation of ERα transcriptional coregulator complexes in the absence of 17β-estradiol (E2), a requirement of WT ERα [17–19]. Y537S is perhaps the most clinically relevant because breast cancer cells harboring the mutant are more metastatic and resistant to first and second-line hormone therapies [20, 21]. Initial studies suggested that SERD (ERα-degrading) activity was required to achieve improved efficacy in Y537S *ESR1* breast cancer cells [22, 23]. However, we recently evaluated a panel of SERMs and SERDs and showed that ERα-degrading activities did not correlate with antagonistic efficacy in this setting [24]. Rather, the most effective SERMs and SERDs favored the formation of a new S537-E380 hydrogen bond that stabilized the LBD antagonist conformation. This interaction is sterically disallowed in the WT Y537 ERα LBD.

Our laboratory recently developed a novel isoquinoline-based SERM, T6I-29, based on structural insights from the recently approved elacestrant and other SERMs and SERDs, to better understand mechanisms of hormone therapy efficacy in Y537S *ESR1* breast tumors [25]. The active enantiomer, T6I-29-1A, showed significant anti-proliferative activities in cultured ER+ breast cancer cell lines; however, its anti-tumoral activities remained to be examined *in vivo* [25]. In this paper, we reveal how T6I-29 interacts with Y537S ERα LBD to engage anti-proliferative activities, downregulate target genes, and elicit anti-tumoral activities *in vivo*. Importantly, we identify neomorphic antiestrogenic activities through the downregulation of DKK1, a tumor-secreted glycoprotein that is associated with metastasis in other cancers [26–29]. Subsequent profiling of circulating DKK1 shows a significant elevation of DKK1 in the plasma of ER+ breast cancer patients versus healthy controls, which increases with tumor stage.

## RESULTS

### T6I-29 Enforces the Antagonist Conformation of the Y537S ERα Ligand Binding Domain

The T6I SERM scaffold adopts a unique ligand binding pose within the WT ERα hormone binding pocket to favor the therapeutic ligand binding domain (LBD) helix 12 (H12) antagonist conformation [25]. It also shows effective anti-proliferative activities in Y537S *ESR1* MCF7 breast cancer cells [25]. Here, we solved an x-ray co-crystal structure of T6I-29 in complex with Y537S ERα LBD to reveal the structural basis of anti-cancer activities. The T6I-29 structure was solved to 2.20 Å with a canonical ERα homodimer in the asymmetric unit. **Figure 1** shows the structural analysis of the Y537S ERα LBD-T6I-29 complex. **Figure 1A** shows an overview of the Y537S ERα LBD homodimer-T6I-29 complex. In the “B” monomer, there are significant crystal contacts in the H11-12 loop and H12 regions confounding analysis. Therefore, analysis is primarily based on the “A” monomer where these crystal contacts are not present.

**Figure 1:**
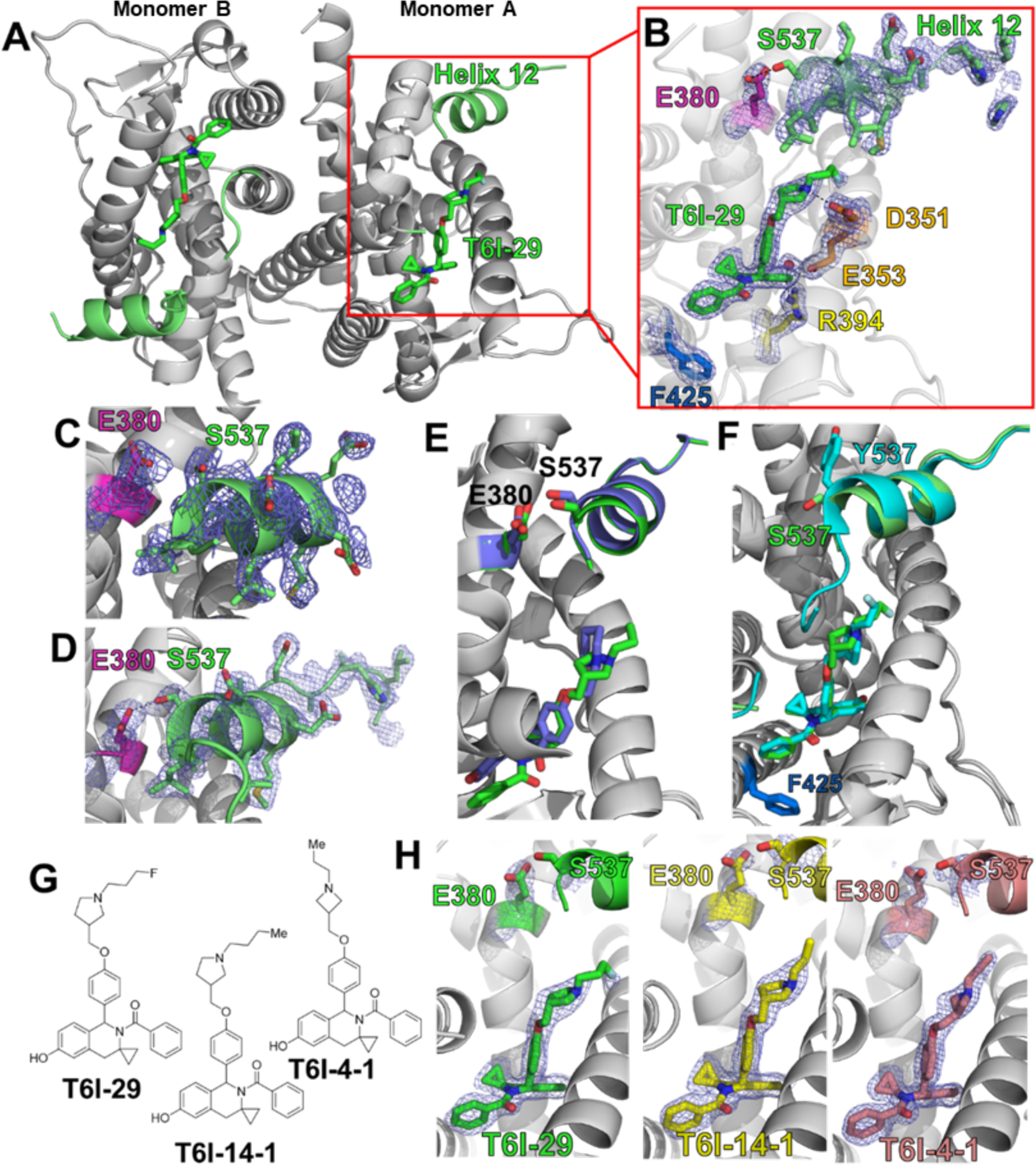
Structural basis of T6I-29 efficacy in Y537S *ESR1* breast cancer cells. A) Overview of the Y537S ERα LBD homodimer x-ray co-crystal structure with T6I-29 (green sticks) bound in the hormone binding pocket. Helix 12 (H12) is highlighted in green. B) T6I-29 interactions with residues in the hormone binding pocket, the difference density map of relevant atoms are shown in blue mesh. C) Difference density map from the Y537S-4OHT x-ray co-crystal structure highlighting the poor density of H12 that is representative of poor transcriptional antagonists in Y537S *ESR1* breast cancer cells. D) Difference density map from the Y537S-RAL x-ray co-crystal structure highlighting the improved density of H12 that is found in effective transcriptional antagonists in Y537S *ESR1* breast cancer cells. Superposition of T6I-29 (green) with RAL (blue) x-ray co-crystal structures. F) Superposition of T6I-29 in complex with WT (cyan) or Y537S (green) ERα LBD. G) Chemical structures of T6I-29, T6I-14-1, and T6I-4-1. H) Side-by-side comparison of ligand, E380, and S537 difference density maps for T6I-29, T6I-14-1, or T6I-4-1 in complex with Y537S ERα LBD. All difference density maps are 2mFo-DFc and are contoured to 1.0 σ. Protein DataBank (PDB) accession codes are: 9BPX for Y537S-T6I-29, 7UJ8 for Y537S-4OHT, 7UJC for Y537S-RAL, 8DVB for WT-T6I-29, 9BQE for Y537S-T6I-14-1, and 9BU1 for Y537S-T6I-4-1.

T6I-29 is resolved in the hormone binding pocket, but reduced difference density is observed in the fluoropropyl group suggesting that the side-arm is more mobile in the Y537S than the previously described WT LBD (**Figure 1B**) [25]. The isoquinoline core forms a hydrogen bond network with E353, R394, and a water molecule within the hormone binding pocket, while the pyrrolidine side-arm forms a hydrogen bond with D351 and the fluorpropyl group adopts a conformation between D351 and helix 12 (H12) (**Figure 1B**). Our earlier study showed that the ineffective SERM 4-hydroxytamoxifen (4OHT) poorly enforced the Y537S H12 antagonist conformation with S537 at too great a distance to form a hydrogen bond with E380 (**Figure 1C**) [19], whereas effective molecules like raloxifene (RAL) maintained a WT-like antagonist conformation with a well resolved H12 and a hydrogen bond between S537 and E380 (**Figure 1D**). Compared to existing structures of SERMs and SERDs in complex with Y537S ERα LBD, the T6I-29-bound structure is most like raloxifene (RAL), which showed significant anti-transcriptional efficacy in breast cancer cells harboring Y537S *ESR1* [24]. H12 is superimposable between the RAL and T6I-29 structures. However, the 537S side chain is poorly resolved in the T6I-29 structure (**Figure 1B**) suggesting that, while more effective than 4OHT, it is less effective than RAL.

Effective SERMs and SERDs maintain a WT-like H12 antagonist conformation when Y537S mutation is present [24]. Here, few differences are observed between the WT and Y537S T6I-29 x-ray co-crystal structures (**Figure 1F**). H12 in the Y537S structure lies in a slightly altered position but is still docked in the AF-2 cleft compared to the WT. This suggests that there is only a minor impact to the H12 antagonist conformation due to the presence of the mutation. Interestingly, the unique impact of T6I-29 on F425 conformation is maintained between the WT and Y537S ERα LBD co-crystal structures (**Figure 1F**). We also solved x-ray crystal structures of analogous T6I-SERMs T6I-14-1 and T6I-4-1 to better understand the structural-basis of activities. The T6I-14-1 structure was solved to 1.98 Å, and the T6I-4-1 structure was solved to 1.75 Å. Compared to T6I-29, T6I-14-1 lacks a fluoro group on the propyl side arm while T6I-4-1 contains a propylazetidine size arm (**Figure 1G**). In each case the T6I core adopts an identical conformation and few conformational differences are observed in H12, S537, and E380 (**Figure 1H**). Therefore, different side-arms can be accommodated on the T6I scaffold to induce the effective H12 conformation in Y537S ERα LBD.

The Y537S ERα LBD mutation can impact the conformational dynamics of the SERM or SERD-saturated complex [19, 30]. Atomistic molecular dynamics simulations were performed to identify potential differences in the mobility between WT and Y537S ERα LBD in complex with 4OHT, lasofoxifene (Laso), T6I-29, or elacestrant (Rad1901). 4OHT is a major active metabolite of tamoxifen and is a SERM that shows reduced efficacy in the presence of *ESR1* LBD mutations [19, 20]. Laso is also a SERM, but it retains efficacy in the presence of Y537S ERα [31]. It is currently in clinical trials (ELAINE trials) for treatment of advanced stage *ESR1* mutant breast cancer [31, 32]. Rad1901 has recently been approved for treatment of advanced *ESR1* mutant breast cancers [33, 34]. In all the simulated systems, the root mean squared fluctuation (RMSF) is low except in regions with the residues 322-342, 392-422, 452-472, and 522-535 (H11-12 loop). Differences in the molecular dynamics induced by the Y537S mutation were most pronounced in the H11-12 loop (residues 525-536) (**Supplemental Figure 1**). For each complex, the Y537S mutant has a much higher RMSF than WT in the H11-12 loop region. These higher fluctuations are consistent with the poorly resolved electron density of the x-ray crystal structures. Interestingly, T6I-29 appears to increase the RMSF to the greatest extent of any of the ligands in the WT LBD, suggesting that it may have unique effects on this region of the protein.

### T6I-29-1A Attenuates the Proliferation, Migration, and ERα Target Gene Upregulation in Breast Cancer Cells Harboring Y537S ESR1

The active enantiomer of T6I-29, T6I-29-1A, was first assessed for its anti-proliferative activities in Y537S *ESR1* breast cancer cell lines compared to clinically relevant compounds and other T6I SERMs. Clinically relevant compounds included fulvestrant (ICI), 4OHT, Laso, Rad1901, and giredestrant (Gir). Rad1901, an orally available SERD also retains efficacy in the presence of *ESR1* mutations, and was recently FDA-approved for patients with *ESR1* mutated advanced ER+ breast cancer based on the positive results of the phase III EMERALD trial [35]. Gir is an orally available SERD, also in clinical trials for treatment of advanced ER+ breast cancers [36, 37]. **Figure 2** shows the impact of T6I-29-1A on Y537S *ESR1* cell proliferation and ER target gene regulation. To assess anti-proliferative effects, T47D Y537S *ESR1* and MCF7 Y537S *ESR1* breast cancer cells were treated with 1 μM compound in the presence of 1 nM estradiol (E2) and changes in cell count were measured over time (**Figure 2A/B**). In both cell lines, T6I-29-1A significantly blunts cell proliferation comparable to other clinically relevant compounds (**Figure 2A/B**). Other T6I compounds T6I-4-1, T6I-6-1, and T6I-10-1 showed limited success in blunting proliferation in both *ESR1* mutant cell lines (**Figure 2A/B**).

**Figure 2:**
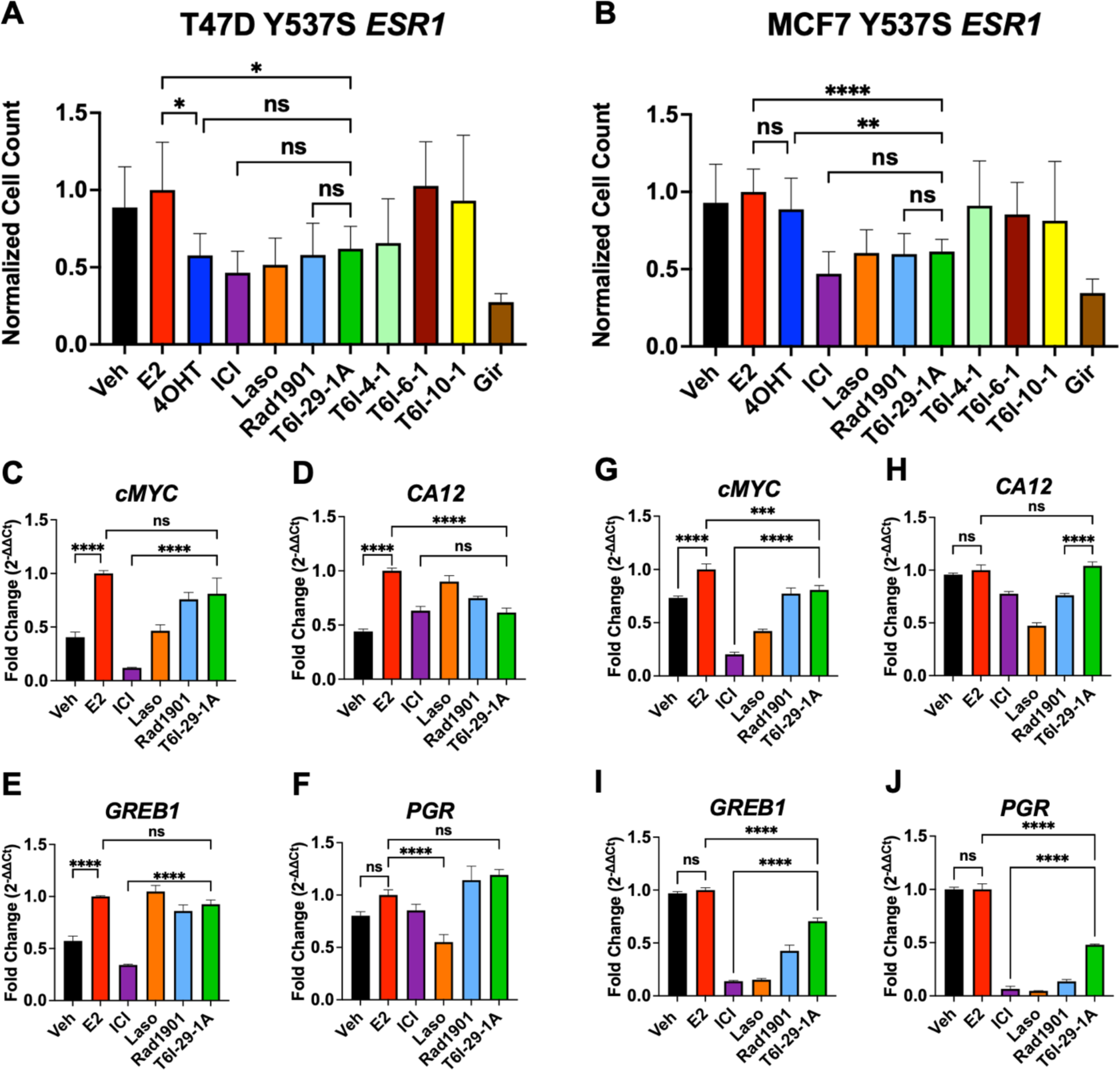
The impact of T6I-29-1A on the proliferation and ER target gene expression of Y537S *ESR1* mutant breast cancer cells. A) T47D Y537S *ESR1* and B) MCF7 Y537S *ESR1* breast cancer cell proliferation, treated with 1 μM compound in the presence of 1 nM E2. Graphs represent mean of three independent replicates, data normalized to E2 treatment, error bars are s.d. Statistical analysis was performed using ANOVA with Tukey post-hoc test. C-F) RT-qPCR in T47D Y537S *ESR1* and G-J) MCF7 Y537S *ESR1* cells. Representative data are the mean of three replicates ± s.d. and error bars show s.d. Significance determined by one-way ANOVA test with tukey post-hoc where *p < 0.05, **p < 0.005, ***p < 0.0005, and ****p < 0.00005.

We next measured the ability of T6I-29-1A to inhibit migratory and stem cell phenotypes of MCF7 Y537S *ESR1* cells. A scratch wound assay showed T6I-29-1A significantly blunted migration in Y537S *ESR1* MCF7 cells (**Supplemental Figure 2A-C**). Mammosphere assays, or 3D colony formation assays, assess the “stemness” of the breast cancer cells [38, 39]. T6I-29-1A decreased the size of mammospheres, but not the total number compared to control, while other relevant compounds decreased both the size and number (**Supplemental Figure 3A-C**).

To investigate effects of T6I-29-1A on ER target gene regulation, we performed RT-qPCR on both MCF7 and T47D Y537S *ESR1* mutant cell lines. Cells were treated with 1 μM compound in the presence of 1 nM E2. In Y537S *ESR1* T47D cells, T6I-29-1A potently downregulated the ER target gene *CA12*, but did not significantly decrease expression of *PGR*, *GREB1,* and *cMYC* (**Figure 2C-F**). Conversely, in Y537S *ESR1* MCF7 cells, T6I-29-1A significantly downregulated ER target genes *GREB1*, *PGR*, and *cMYC*, but did not *CA12* (**Figure 2G-J**). Although it appears that T6I-29 does not downregulate ERα target genes as potently as ICI (with the exception of *CA12* in T47D Y537S *ESR1* cells) it behaves similarly to Rad1901, recently approved for *ESR1* mutated advanced metastatic breast cancer.

### Inhibition ERα-Coactivator Binding

In ER+ breast cancers, ERα recruits various coactivators to fuel transcriptional-driven tumor growth, with steroid receptor coactivator-3 (SRC3) being one of the most associated with pro-oncogenic activities [40–42]. SERMs and SERDs favor a H12 conformation that disfavors SRC3 binding via LXXLL motifs in the activating function-2 cleft of ERα [43]. We used the NanoBiT assay to measure how T6Is and other relevant compounds impacted the association between SRC3 and WT or Y537S ERα [44]. Clinically relevant compounds used in this assay included: Gir, ICI, amcenestrant (Amc), camizestrant (Cam), 4OHT, Laso, Rad1901, and lead T6Is (T6I-29-1A, T6I-4-1, and T6I-6-1). The SERD Amc was recently discontinued after phase II clinical trials after failure to meet primary endpoints [45]. Cam is an oral SERD currently in clinical trials [46, 47]. Plasmids encoding either wild-type (WT) smBiT-ERα or mutant smBiT-Y537S ERα were co-transfected with a plasmid encoding lrgBiT-SRC3 into HEK293T cells. Following transfection, cells were introduced into charcoal-stripped serum depleted of hormone for 72 hours. Cells were then treated with serial dilutions of SERM or SERD (5-fold from 5 μM to 12.8 pM in triplicate, over three biological replicates) in the presence of 1 nM E2. Each plate included DMSO and 1 nM E2 control wells in triplicate. After 48 hours of treatment, which showed the best signal-to-noise ratio, wells were read for luminescence. From this, we derived IC50 data for WT-SRC3 and mutant Y537S-SRC3 interactions in the presence of different drug treatments. **Figure 3** shows the IC50s of relevant clinical compounds and T6Is on this protein-protein interaction.

**Figure 3:**
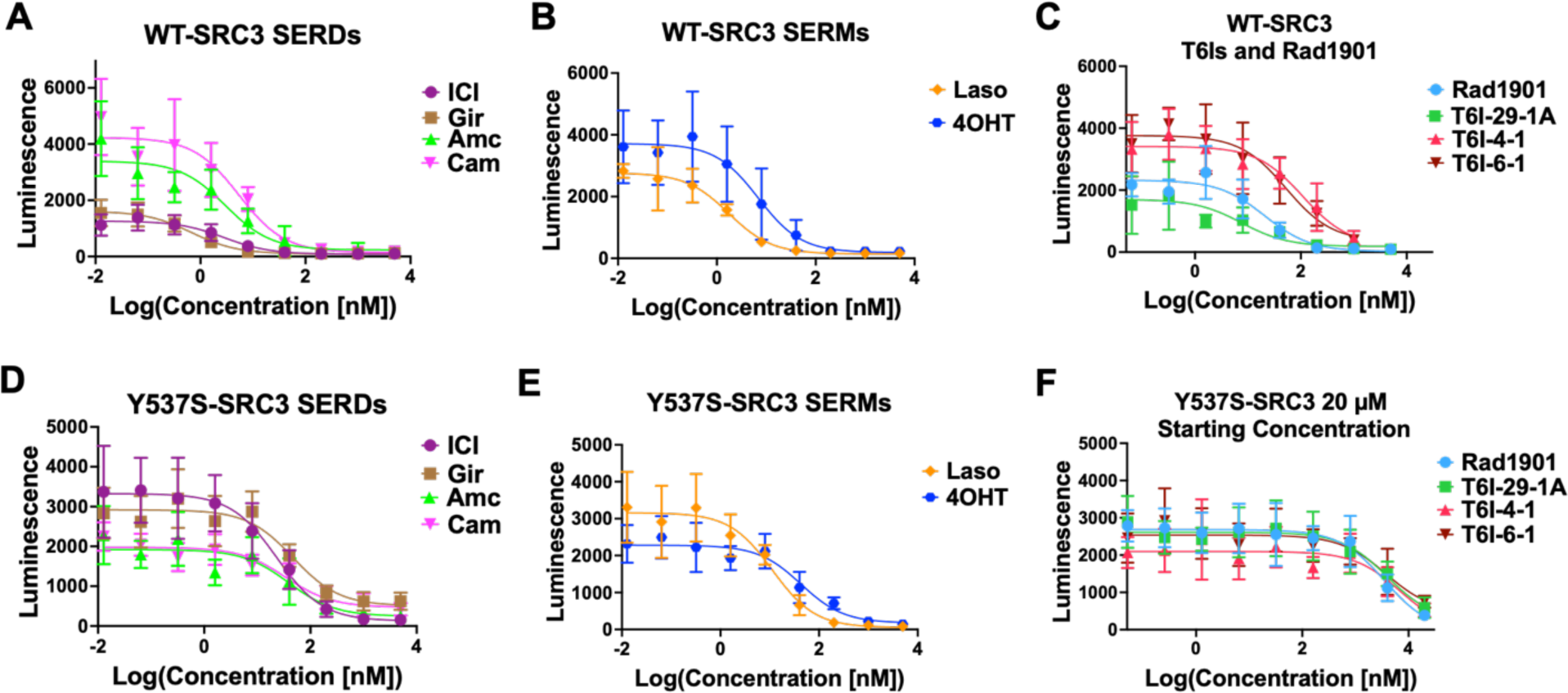
Lead T6Is and clinically relevant SERMs and SERDs inhibit WT and Y537S ERα-Coactivator binding. Clinically relevant A) SERD and B) SERM inhibition curves with WT-SRC3 binding. C) Rad1901 and T6Is inhibition curves for WT-SRC3 binding. D-F) Same as A-C, but with Y537S-SRC3 binding. Data are shown as the mean ± s.d.

**Table 1** shows IC50 values for each compound in WT and Y537S-SRC3 interactions. All compounds tested showed increased inhibitory potency in the WT setting compared to Y537S **(Table 1)**. In the WT setting, SERDs including Gir and ICI demonstrated the greatest potency followed by SERMs Laso and 4OHT, while Rad1901 and the T6Is, including T6I-29-1A, had the lowest IC50s **(Figure 3A-C**, **Table 1)**. In the Y537S setting, Laso showed the greatest inhibitory potency while Rad1901 and the T6Is remained the least potent **(Figure 3D-F**, **Table 1)**. It should not be surprising that Laso showed the greatest potency in the presence of the mutant since it also maintains its binding affinity and enforcement of the LBD antagonist conformation [31]. Both Rad1901 and the T6Is required additional treatments up to 20 μM in order to measaure IC50 values in the Y537S setting **(Figure 3F)**. In concordance with these findings, there is a larger difference in IC50 values between WT and Y537S in SERMs 4OHT, Rad1901 and the T6I compounds compared to SERDs (ICI, Gir, Amc, Cam) **(Table 1)**. Based on these data, Rad1901 as well as the T6Is may primarily function to blunt tumor growth via other mechanisms of antagonism than this specific coactivator interaction with ERα and SRC3.

**Table 1:**
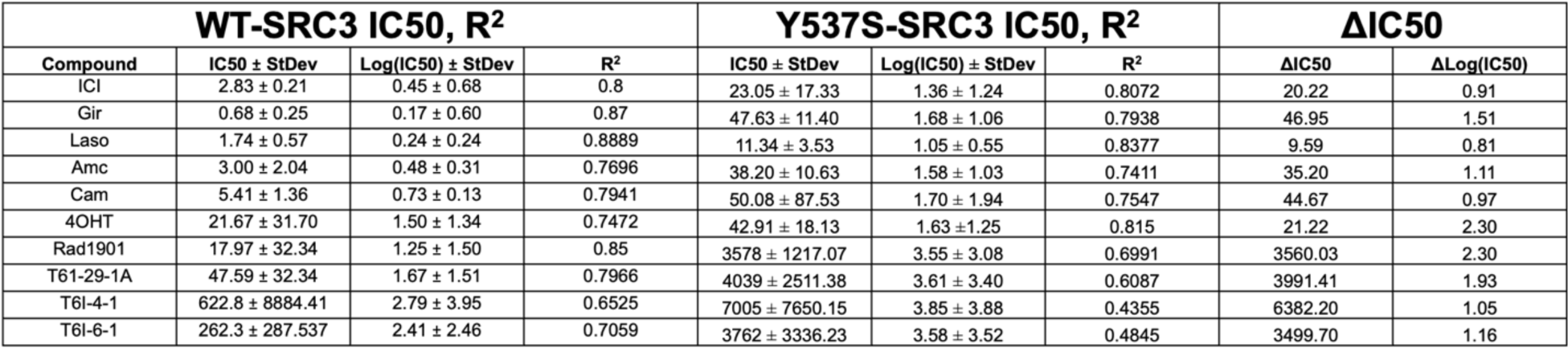
IC50s and standard deviations of clinically relevant and T6I compounds on inhibition of receptor-coactivator interaction. Left: IC50s, standard deviation, and R^2^ of WT-SRC3 co-transfection interaction. Middle: IC50s, standard deviation, and R^2^ of Y537S-SRC3 co-transfection interaction. Right: Differences in IC50 values between WT-SRC3 and Y573S-SRC3 co-transfection interactions. All data represents three biological replicates.

### Pharmaceutical Properties of T6I-29

Preliminary drug metabolism and pharmacokinetics (DMPK) and adsorption, distribution, metabolism (ADME) were measured to determine the suitability of T6I-29 for *in vivo* studies. **Figure 4** shows DMPK and ADME profiles of T6I-29 *in vivo* preliminary studies. For the DMPK studies, 25 mg/kg was chosen as the starting dose and it was tested by intraperitoneal (IP) and oral gavage (PO) administration routes in C57/BL/6J mice (**Figure 4 A/B**). For drug delivery vehicle we used 20% DMSO dissolved in 20% captisol in water for IP and the pH was adjusted with HCl. For PO, 2% tween 80 and 0.5% methylcellulose was used in water (**Supplemental Tables 1/2).** T6I-29 shows a serum half-life is 3.60 ± 0.07 and 4.02 ± 0.96 hours by IP and PO respectively. Its mean Cmax was 5,053 ± 995 and 752 ± 70 ng/mg for IP and PO respectively. The AUC was 8,350 ± 1,038 and 2,931 ± 503 h*ng/mL for IP and PO respectively. The ADME for T6I-29 in human and mouse plasma protein binding showed that 1.54% and 2.57% fraction unbound by protein respectively. This ADME profile is similar to other SERMs and SERDs, with tamoxifen also showing greater than 98% protein binding [48]. No signs of toxicities were observed in these preliminary studies.

**Figure 4:**
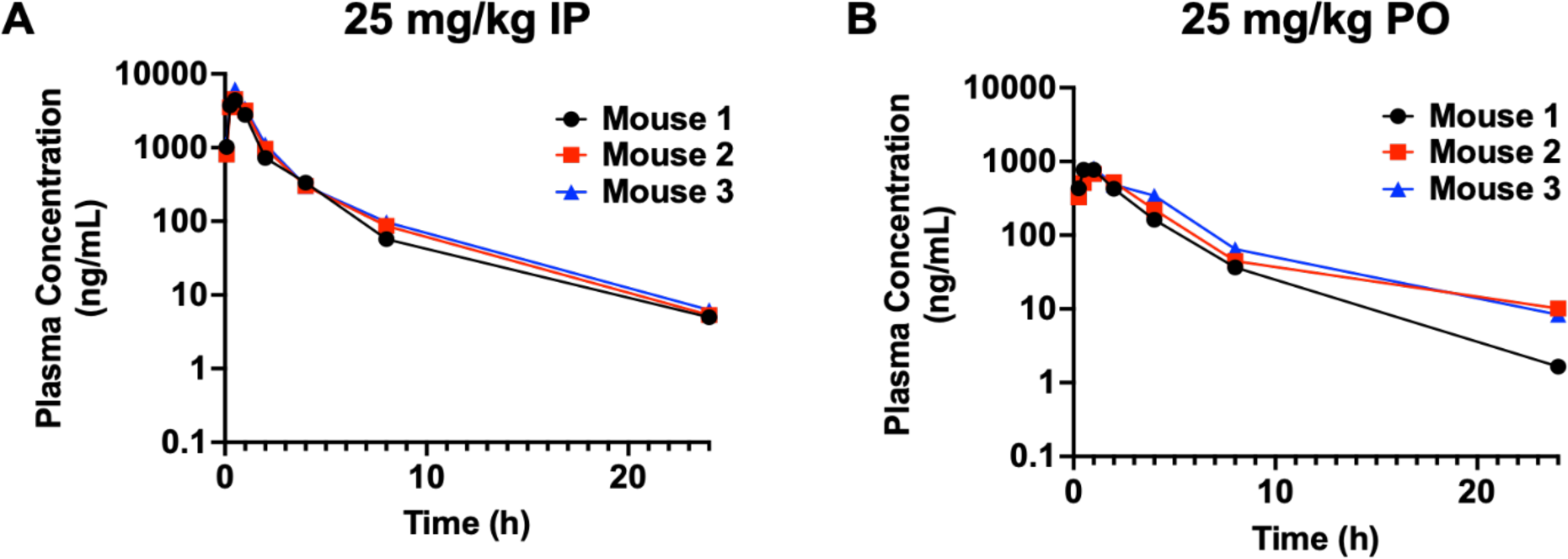
Pharmacokinetics of T6I-29-1A measured at 25 mg/kg dose A) By IP and B) By PO. Serum half-life was interpolated from curves. Three mice were used per study. Plasma concentration measured by ELISA.

### Pilot Murine T6I-29-1A Studies

To characterize the effects of T6I-29 on tumor growth and to determine the best mode of delivery, we used an ectopic murine Y537S *ESR1* MCF7 xenograft model and treated with different doses of T6I-29-1A. Female NOD/SCID ovariectomized mice were bilaterally injected with homozygous Y537S *ESR1* MCF7 cells in their mammary fat pads. After tumors reached 100 mm^3^, mice were randomized into different treatment groups. **Figure 5** shows anti-tumoral effects of T6I-29-1A in preliminary *in vivo* IP and PO studies. We found that via IP injection five times a week, T6I-29-1A appeared to significantly inhibit tumor growth at 25 mg/kg and 100 mg/kg doses, measured at day 9 of treatment (n= 4-8 tumors/treatment group) as measured by caliper three times per week (**Figure 5A**). Examining metastatic lesions at common sites (liver, lung, brain, femurs, and uterus) by pathologist Dr. Khin Su Mon showed the fewest number of metastases occurred with the 25 mg/kg dose of T6I-29-1A (**Figure 5B**). There was no significant uterine stimulatory or antagonistic effects with any dose of T6I-29-1A (**Supplemental Figure 4A/B**). We did not observe a significant survival benefit with any dose of T6I-29-1A by IP in this pilot study but the 100 mg/kg cohort trended towards significance (*p* = 0.056). (**Figure 5C**). Representative metastatic lesions in the liver, adrenal gland, femur, and uterus by H&E stain are shown (**Figure 5D**).

**Figure 5:**
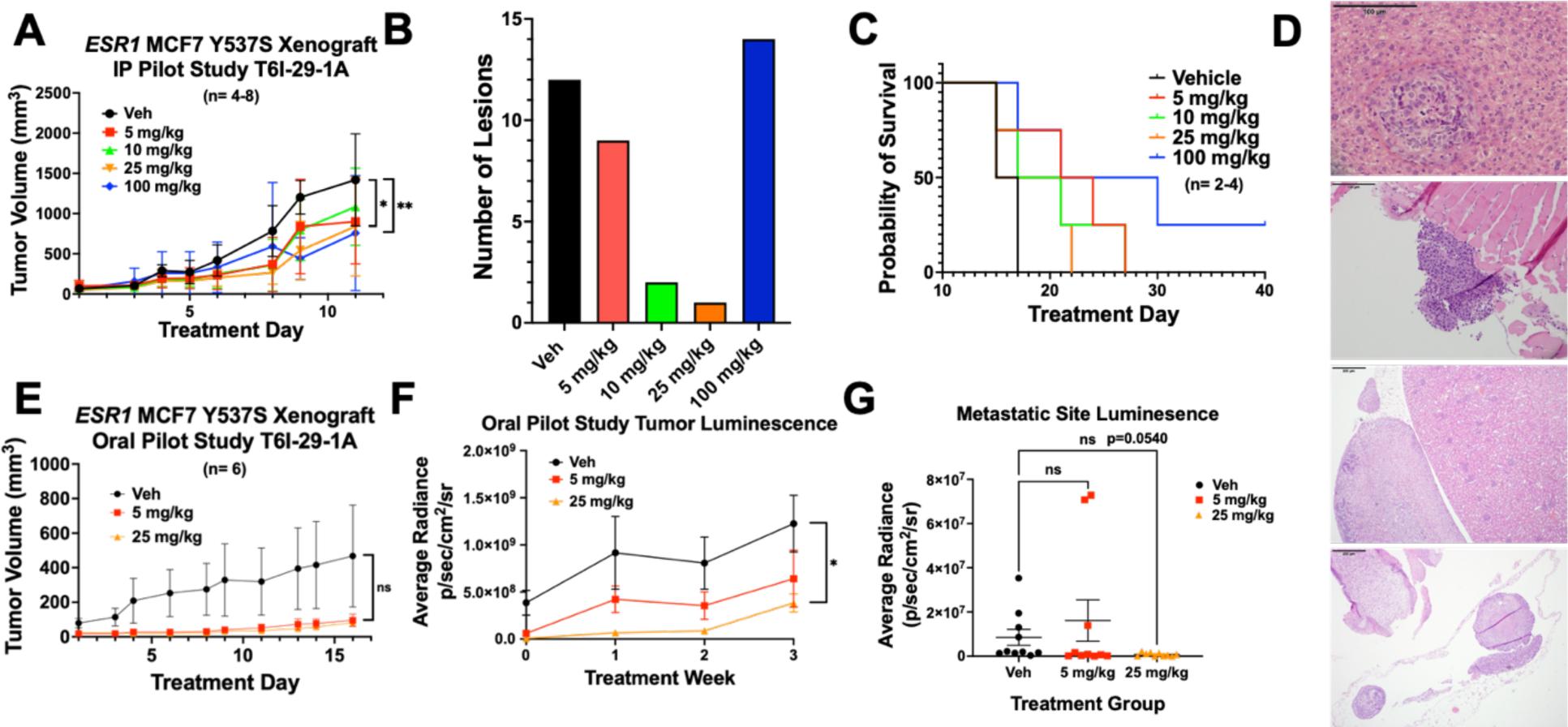
T6I-29-1A inhibits tumor growth in preliminary *in vivo* studies. A) Tumor growth (error bars indicate SEM) in I.P. pilot study, n= 4-8 tumors/ group. Significance is measured by Two-Way Anova with Bonferroni post-hoc test, results indicate day 9 treatment analysis. B) Total metastatic lesions as measured by H&E staining by Dr. Khin Su Mon across groups. C) Survival curve of I.P. pilot study, significance determined using log rank test. Veh vs 100 mg/kg p=0.0624. D) Representative photos capturing metastases (top to bottom) in liver (vehicle treated), left femur (vehicle treated), adrenal gland (100 mg/kg treated), and uterus (vehicle treated). E) Tumor growth (error bars indicate SEM) in oral pilot study, n= 6 tumors/ group. Analyzed with Two-Way Anova with Bonferroni post-hoc test. F) Tumor luminescence of oral pilot study measured weekly (error bars indicate SEM). Analyzed with unpaired t-test at treatment week 3. G) Luminesence of liver, lung, brain, femurs, uterus were measured for each mouse in each group *ex vivo* (error bars indicate s.d.), results were graphed based on treatment groups, including both sides of organ luminescent signal. Anova with Tukey post-hoc statistical test was used to determine significance.

To investigate whether oral administration maintained tumor blunting activities, heterozygous, luciferase tagged Y537S *ESR1* MCF7 were used. Using the same cell injection and mouse randomization protocol, we monitored tumor growth with 5 and 25 mg/kg doses of T6I-29-1A, administered five times per week by oral gavage. By caliper, T6I-29-1A did not appear to significantly inhibit tumor growth (**Figure 5E**). However, tumors were also analyzed using bioluminescence imaging with the IVIS system, and through this method, T6I-29-1A significantly diminished tumor growth compared to vehicle at 25 mg/kg treatment (**Figure 5F**). *Ex vivo* analysis of common sites of breast cancer metastasis (liver, brain, femurs, uterus) showed a trend toward significant decrease with increasing dose of T6I-29-1A (**Figure 5G**). To this end, metastatic characterization by Dr. Marteen Bosland confirmed some metastatic lesions as determined by IVIS system (**Supplemental Figure 5A-C**). However, very few metastatic lesions were found overall via histological staining, revealing shortcomings of this xenograft model. We did not observe any uterine stimulatory or degradation with oral dosing of T6I-29-1A (**Supplemental Figure 6C/D**).

### ICI Exhibits Improved Tumor Growth Inhibition Compared to T6I-29-1A

Based on our preliminary murine pilot IP and PO studies (**Figure 5**), we used a 25 mg/kg IP dose in a comparative study to ICI to investigate tumor growth and metastatic colonization differences between treatment conditions with increased statistical power. Using the same ectopic xenograft model, heterozygous luciferase tagged Y537S *ESR1* MCF7 cells were bilaterally injected into the mammary fat pads of female NOD/SCID ovariectomized mice, and mice were randomized to different treatment groups when tumors reached 100 mm^3^ (10 mice/ group). **Figure 6** shows anti-tumoral effects of T6I-29-1A compared to a clinical standard for advanced ER+ breast cancer, ICI. Mice were treated with Vehicle (Veh), 25 mg/kg T6I-29-1A five times per week, or a clinically relevant dose of 25 mg/kg ICI once per week [49]. We observed a reduced but not significant reduction in tumor growth in the T6I-29-1A-treated group compared to vehicle, while ICI significantly blunted tumor growth, as measured by digital caliper **(Figure 6A, Supplemental Figure 7A).** While ICI significantly decreased final uterine weights, T6I-29-1A had no significant stimulatory or degrading effects **(Figure 6B)**. Previous studies have shown that SERM treatment increases endometrial thickness due to estrogenic nature of compounds, while SERDs such as ICI, inhibit growth [50–52]. To this end, rodent uterine models are used to assess estrogenic-stimulatory capacity of compounds, and higher estrogenic stimulation may indicate higher risk for endometrial cancer [49, 53, 54]. In an additional uterine SERM agonist study, we measured endometrium thickness in female BALB/c ovariectomized mice treated with 4OHT, ICI, and T6I-29-1A in the presence and absence of E2 compared to vehicle with and without E2 treatment **(Supplemental Figure 8)**. Based on analysis of endometrium thickness, 4OHT treatment significantly increased width, T6I-29-1A treatment did not have a significant effect, and ICI diminished the thickness **(Supplemental Figure 8A-E**).

**Figure 6:**
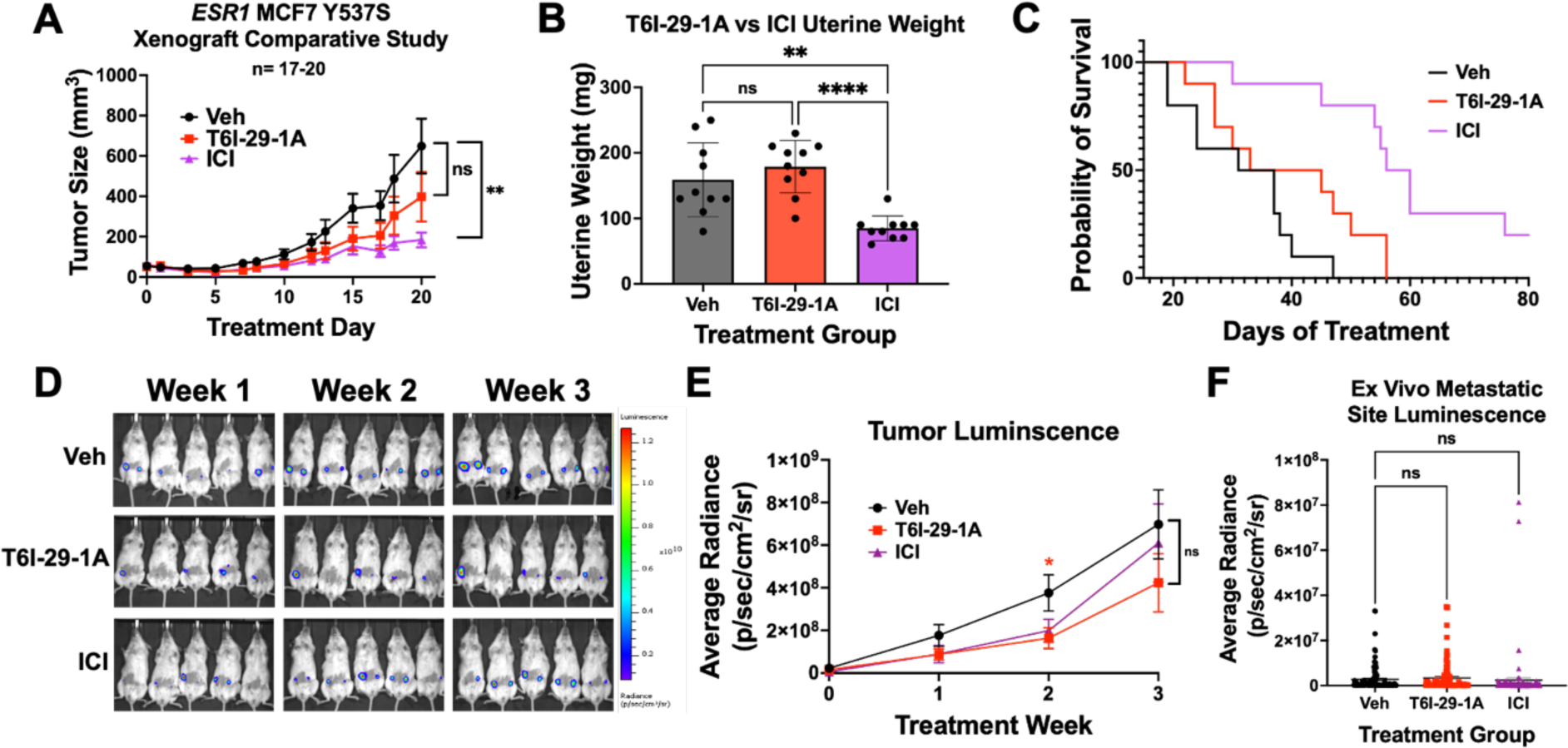
ICI blunts tumor growth more effectively than T6I-29-1A. A) Tumor growth (error bars represent SEM) in vehicle, T6I-29-1A, and ICI treatment groups (n=17-20 tumors/ group). Significance is measured by Two-Way Anova with Bonferroni post-hoc test. B) Final uterine weights (n= 10 mice/ group). Significance is measured by Anova with Tukey post-hoc test. C) Survival curve compared to vehicle. Log Rank test used to determine significant survival benefit. Veh vs. T6I-29-1A: p=0.0966, Veh vs. ICI: p=0.0001, T6I-29-1A vs. ICI p=0.0052. D) Representative weekly IVIS bioluminescent imaging denoting weekly tumor growth. Scale bar shown on right. E) Quantified luminescence for each treatment group (error bars represent SEM). Significance was determined using unpaired t test at each week with Mann-Whitney correction. F) Ex vivo metastatic luminescence at common sites (liver, femurs, uterus, brain) for each mouse. Anova with Tukey post-hoc was used to determine significance.

In the comparative study with T6I-29-1A and ICI, survival increase was not significant for mice treated with T6I-29-1A (p= 0.0966), while it was significantly prolonged for ICI treated mice (**Figure 6C**). Tumor growth was also monitored via bioluminescent imaging using the IVIS system. We observed that tumor luminescence signal was significantly diminished by T6I-29-1A, measured at treatment week 2, but tumor luminescence was non-significant at week 3, indicating a potential early anti-tumor effect that is lost over time (**Figure 6E**). However, there was no significant difference in bioluminescent signal when comparing ICI to vehicle (**Figure 6E**). IVIS *ex vivo* analysis showed no statistical difference in metastatic bioluminescence of common sites (liver, brain, femurs, uterus) (**Figure 6F**). While some of these metastatic sites measured by IVIS were confirmed by pathology analysis of H&E stained tissues, these results showed very little metastatic burden across any group (**Supplemental Figure 9A-E**). Individual sites metastatic luminescence was quantified individually, all with no significant change in metastases, with exception of right femur **(Supplemental Figure 10A-G**). RT-qPCR was used to quantitate ER target gene effects with different treatment groups, with trends towards downregulation in T6I-29-1A treated mice, that is heightened with treatment of ICI, although no significance was noted **(Supplemental Figure 11A-G).**

### T6I-29 Uniquely Downregulates DKK1 in Y537S ESR1 Breast Cancer Cells

Structurally unconventional SERMs and SERDs can reveal new ER-coregulator interactions and transcriptional activities [25, 55]. In WT *ESR1* breast cancer cells T6I-29-1A showed unique effects on genes related to SUMO and SUMOylation [25]. Here, RNA-sequencing was used to determine whether T6I-29-1A engaged unique transcriptional programs in MCF7 Y537S *ESR1* cells. RNA was isolated from cells treated with relevant clinical compounds (ICI, Laso, and Rad1901) and T6I-29-1A at 1 μM in the presence of 1 nM E2 for 24 hours. **Figure 7** shows distinct transcriptional programs engaged by T6I-29-1A compared to other SERMs and SERDs. While there was significant overlap between all treatment conditions, T6I-29-1A uniquely and significantly downregulated pathways associated with cell morphogenesis and components of the extracellular matrix (**Figure 7A/B**). As we previously observed in WT *ESR1* cells T6I-29-1A shares the most differentially expressed transcripts in common with ICI (**Figure 7C**) [25]. Pathway analysis showed that T6I-29-1A uniquely impacted genes associated with the Wnt/β-Catenin pathway, including cell adhesion and morphogenesis (**Figure 7D**).

**Figure 7:**
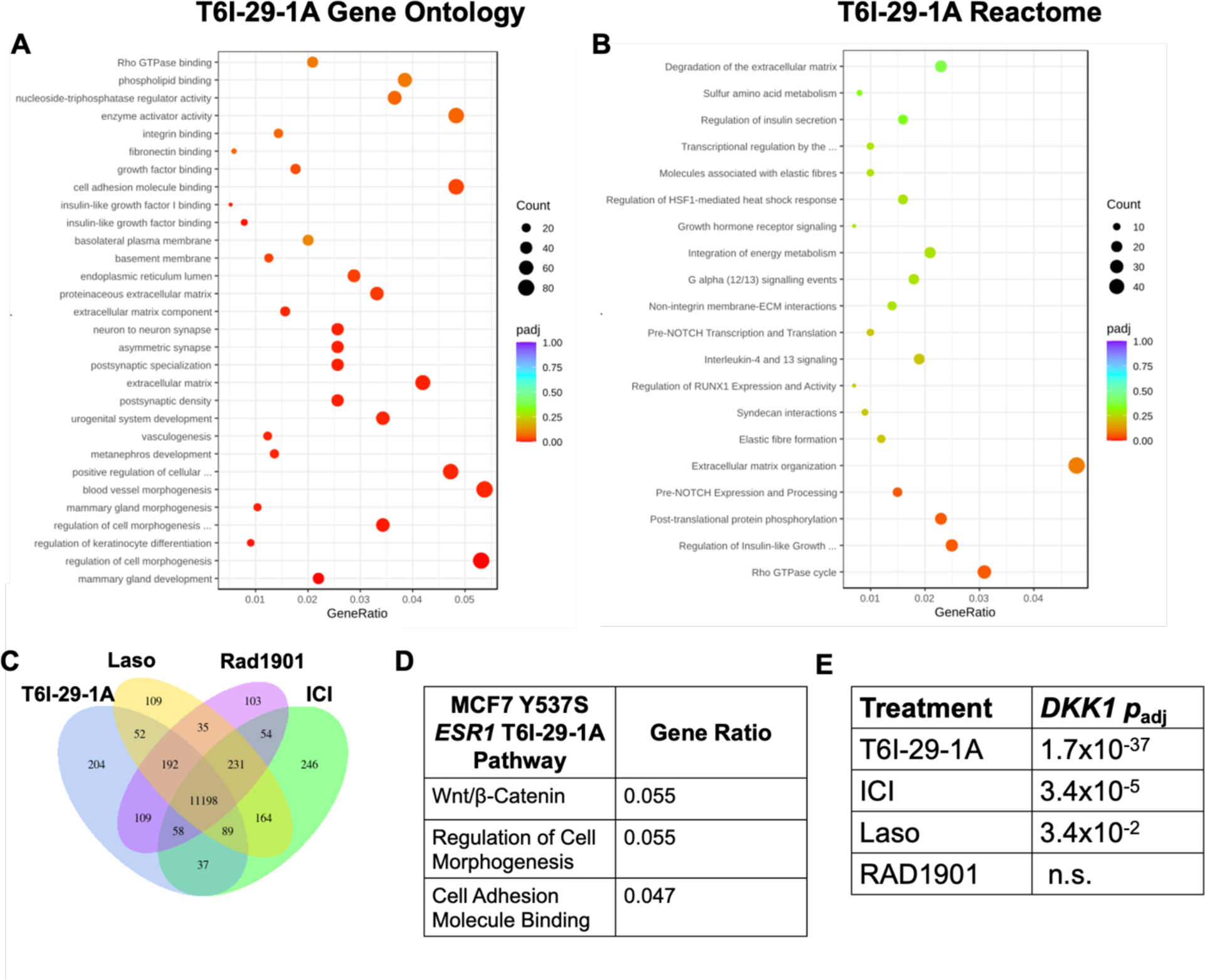
RNA-sequencing reveals DKK1 downregulation uniquely by T6I-29-1A. A) Gene ontology and B) Reactome of T6I-29-1A. Increasing red color denotes higher significance (smaller P value), with dot size correlating to number of transcripts. GeneRatio refers to number of transcripts changed in T6I-29-1A treated cells versus genes associated with each term. C) Uniquely and shared differentially expressed transcripts with T6I-29-1A, Laso, Rad1901, and ICI. T6I-29-1A uniquely regulates 204 transcripts. D) Pathways most differentially regulated by T6I-29-1A. E) *DKK1* downregulation by different antietrogen treatments. T6I-29-1A most significantly downregulates the gene expression, followed by ICI.

Interrestingly, these were Y537S *ESR1* allele-specific pathways previously shown to enhance the metastasis of breast cancer cells harboring the mutant [56].The gene that was most significantly downregulated by T6I-29-1A is *DKK1*, (gene for Dickkopf-1) a known modulator of the Wnt/β-Catenin pathway (**Figure 7E**) [57]. Based on these findings, we further studied the significance of *DKK1* in ER+ breast cancer.

### DKK1 is Elevated in the Plasma of ER+ Breast Cancer Patients

DKK1 is a secreted glycoprotein that is classically known as an inhibitor in the Wnt/β-Catenin pathway, although it demonstrates non-canonical activities that that are implicated in pathogenic progression across many cancers [26, 57, 58]. In breast cancer, DKK1 is amplified in the serum of breast cancer patients with bone metastases [59, 60]. However, there were relatively few studies to show the patient-relevance of DKK1 expression in ER+ breast cancers. To improve our understanding of the patient-significance of differential DKK1 expression, we profiled DKK1 levels in 108 ER+ breast cancer patient plasma samples compared to 105 matched plasma controls from healthy women. These were obtained from the Simon Cancer Center at Indiana University and the Susan G. Komen Tissue Bank respectively. **Figure 8** shows patient plasma DKK1 concentrations compared to healthy controls. For each sample, an ELISA dilution curve was ran to obtain the linear range of signal absorption (**Supplemental Figure 12A**). Plasma concentrations were interpolated based on a dilution series from recombinant DKK1 adsorbed on each ELISA plate. DKK1 protein levels are significantly higher in ER+ patient plasma compared to healthy controls (**Figure 8A**).

**Figure 8:**
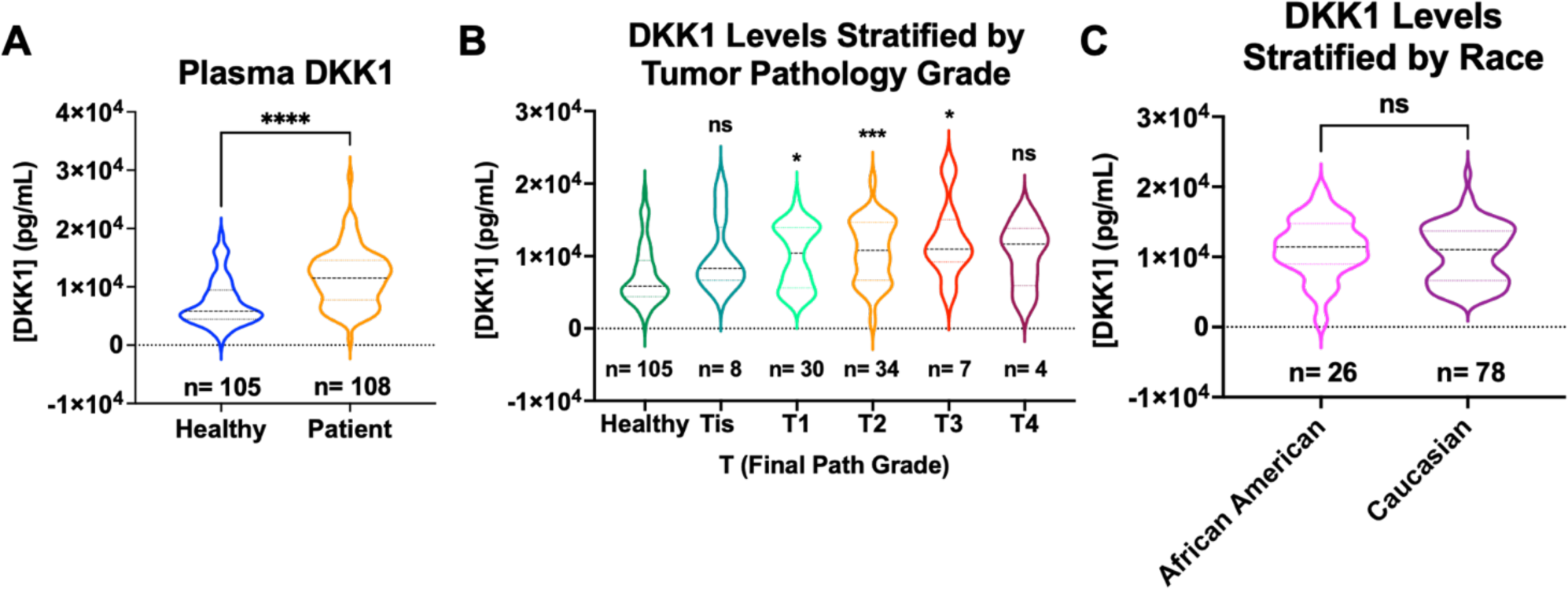
DKK1 levels are increased in ER+ patients compared to healthy control samples. A) Plasma DKK1 values interpolated by ELISA assay, significance determined by unpaired t-test, ****p < 0.00005. B) DKK1 values stratified by T path grade, significance determined by Kruskal Wallace test, with Dunn’s multiple comparison, *p < 0.05, **p < 0.005, and ***p < 0.0005. C) DKK1 values stratified by self-reported race. Significance determined by Mann Whitney test. DKK1 values are all based on a 1 to 100 dilution in the ELISA assay interpolated to the standard curve.

Correlation with available clinicopathologic variables showed that DKK1 levels were elevated in patients with higher pathologic T stage (Tis-T4) with exception of T4 tumors, however this may be due to small sample size (**Figure 8B**). Pathologic T stage relates to primary tumor size, with higher T stage indicative of larger primary tumor size [61]. When stratified by self-reported race, Caucasian patients showed a bi-modal distribution, while African American patients were in the middle of the two distributions but trended towards high levels (**Figure 8C**). So, while DKK1 levels are not significantly different in patients stratified by race, future studies may reveal a disparity in DKK1 expression between these patient populations. Together, these data further add to the body of evidence that DKK1 protein levels are elevated ER+ breast cancer patients compared to healthy controls. They also point to a potential role and ER-dependence in endocrine-resistant breast cancer patients.

## DISCUSSION

In examining the potential utility of the SERM T6I-29-1A in Y537S *ESR1* mutant breast cancers, this study revealed a potential new method to modulate DKK1, a paracine factor associated with metastatic progression in many cancers. Comprehensive structural studies show that T6I-29 engages the S537-E380 hydrogen bond that is associated with improved anti-proliferative efficacy [24]. It also maintains the unique influence on helix 8, specifically perturbing F425, which was observed in the WT ERα LBD co-crystal structure [25]. *In situ* coactivator (SRC3) binding studies show that T6I-29 largely matches the potency and efficacy of elacestrant (Rad1901). Whereby, potency is reduced in the presence of the Y537S mutation, but the interaction can be fully inhibited at higher concentrations.

Pilot *in vivo* studies showed a significant inhibition of tumor growth when Y537S *ESR1* xenograft tumors were treated with T6I-29-1A, with more significant anti-tumoral effects observed in the I.P. pilot study. Interestingly, although T6I-29-1A structurally is more SERM-like, we did not observe any change in uterine weights in any study, even with the highest dosing of T6I-29-1A. This agrees with the reduced alkaline-phosphatase activities that were previously observed in Ishikawa endometrial cells [25]. Therefore, while T6I-29-1A does not induce ERα degradation like a SERM, it does not have the uterine-stimulating liabilities of SERMs like tamoxifen.

Although initially promising, a powered comparative study with ICI (a current clinical standard-of-care) T6I-29-1A falied to significantly decrease tumor burden when measured by digital caliper; however, T6I-29-1A had a significantly diminished luminescent signal compared to vehicle (measured weekly by IVIS Spectrum imaging machine) while ICI did not. When investigating metastatic burden in the mice treated with T6I-29-1A, our studies largely found no effect from the drug versus vehicle treatment. With *ex vivo* measuring of metastatic organs, we observed that T6I-29-1A had a significantly diminished luminescent signal in the oral dosing pilot study in the brain specifically, but overall metastatic burden was not significantly decreased (p=0.0540). However, mice did not show signs of toxicities with one mouse receiving 100 mg/kg daily by I.P. for forty days. It may be the case that optimizing the vehicle formulation, treatment schedule, and increasing the dose will improve the anti-tumoral properties of T6I-29-1A. Moreover, this scaffold will also benefit from further optimization to maximize potency and improve DMPK and ADME.

These *in vivo* studies present additional opportunities for future directions. While we did not see significant differences overall in metastatic lesions, the duration of the studies were quite short, as tumors grew out very quickly. In the future, resecting the initial tumor in the mice, followed by monitioring the mice until tumors reach growth endpoints again might allow for more metastatic lesions to occur and aid in measuring. A limitation of this study is that not many metastatic lesions were found by pathology (H&E) staining, and these MCF7 Y537S *ESR1* cells are known to have metastatic colonization properties [20, 62]. It is also important to note that these tumors are xenograft models, where tumor cells are injected into the mammary fat pad. However, moving forward, other models that better recapitulate the breast microenvironment should be considered. This includes the Mouse Mammary Intraductal Method (MIND) model, which resembles human disease to a greater degree [31, 63, 64].

Unconventional SERMs and SERDs can reveal neomorphic ERα activities by targeting unique LBD structural elements and repopulating coregulator complexes [55]. In WT *ESR1* breast cancer cells, T6I-29-1A uniquely impacted genes related to SUMOylation [25]. In this study, mRNA sequencing shows that T6I-29-1A significantly downregulates DKK1, a paracine factor associated with metastasis in other cancers. The significance of this finding extends beyond cultured breast cancer cell lines because DKK1 is significantly elevated in the blood plasma of ER+ breast cancer patients compared to healthy donor controls. While these findings are exciting, we are limited by the annotation of our initial pilot cohort. As such, further profiling in patients with complete genomic profiling, treatment histories, and outcome is required to understand who these DKK1_high_ patients are. If, as our data suggest, DKK1 expression can be modulated by a SERM and the glycoprotein contributes to metastatic progression, then this study has revealed a new therapeutic axis that can be exploited to treat endocrine-resistant *ESR1* mutant breast cancer.

## MATERIALS AND METHODS

### Chemicals, reagents, and kits

All T6I SERMs were synthesized as previously described [25]. 17β-estradiol was purchased from Millipore Sigma (50-28-2) and used for all experiments. 4-hydroxytamoxifen, fulvestrant, Rad1901, lasofoxifene, giredestrant, amcenestrant, and camizestrant were purchased from MedChem Express (catalog numbers HY-16950, HY-13636, HY-19822, HY-A0037, HY-109176, HY-133017, HY-136255, respectively). All cell culture, bacterial expression media and reagents, and quantitative PCR reagents were purchased from Thermo Fisher Inc. RNA extraction was performed using RNeasy mini kit from Qiagen (catalog number 74106).

### Protein Expression, Purification, and Crystal Structure Determination

Estrogen receptor ligand binding domain (positions 300 – 550) with C381S, C417S, C530S, and Y537S was recombinantly expressed in *E.coli* and purified exactly as described [25]. For the x-ray co-crystal structures with T6Is, 1 mM of each SERM was incubated overnight at 4°C with 400 µM ERα LBD. The next morning, the mixture was centrifuged at 16,000 *xg* for 30 minutes to remove any precipitate. Hanging drop vapor diffusion was used to obtain diffraction quality crystals whereby 2 µL of the LBD mixture at 5 or 10 mg/mL was incubated with 2 µL of mother liquor. After an average of a week rectangular crystals formed in 20-30% PEG 8,000, 200 mM MgCl_2_, and 100 mM HEPES pH 7-8.0. Crystals were cryo-protected in mother liquor with 25% glycerol. X-ray diffraction data sets were collected and processed using the automated protocols at the 17-ID-1 beamline at the NSLS-II, Brookhaven National Laboratories. **Supplemental Table 3** shows the x-ray data collection and refinement statistics. Molecular replacement was used to solve the structure using PDB: 8DUK with the ligand removed as the starting model. Elbow was used to generate ligand constraints. Ligands were placed in the orthosteric hormone binding pocket after clear difference density was observed following the first round of refinement using Phenix refine [65]. Iterative rounds of Phenix refine followed by manual inspection and editing in Coot was used to fully solve the structures. Unresolved atoms were not included in the final model.

### Molecular Dynamics Simulations

Molecular dynamics (MD) simulations were performed of the Estrogen Receptor alpha (ERalpha) wild type (WT) and Y537S mutant monomer, in complexes with four ligands: 4-hydroxytamoxifen (4OHT), lasoxifene (Laso), RAD1901, and T6I-29. Models were built based on the corresponding crystal structures, except that models of the Y537S mutant bound to 4OHT and RAD1901 were based on superposing the ligand into the crystal structure originally solved with T6I-29 (PDB: 9BPX). As the crystal structures contain thermostabilizing mutations of exposed serines to cysteines, these were reverted to wild type using PDBFixer (v1.9). Protein-ligand complexes were protonated at a pH of 7.0 with pdb2pqr30 (v3.6.1) [66]. Amber ff14SB, General Amber force field (GAFF), and OPC3 force fields were used to parameterize protein, ligand, and solvent topologies, respectively [67, 68]. Simulations were performed using the open source MD engine OpenMM (v8.0.0) with the Langevin Middle Integrator maintaining a temperature of 300 K with a timestep of 3 femtoseconds [69]. Constant pressure simulations used a Monte Carlo barostat with a pressure of 1 bar. Each protein-ligand complex was equilibrated with 125 ps of constant volume simulation and 500 ps of constant pressure simulation. Production runs comprising 500 ns of constant pressure simulation for each protein-ligand complex were computed in triplicate. Root-mean-square fluctuations (RMSF) of production simulation were calculated using the open source python package MDAnalysis (v2.4.2) [70, 71].

### Mammosphere Assays (MS)

MCF7 *ESR1* Y537S mutant cells were seeded at single cell density of 400cells/well on 96W low attachment plates. MS medium was prepared according to Dontu et al. and supplemented with 1% methyl cellulose to prevent cellular aggregation [72]. After 7 days in culture, the number and average diameter size of mammospheres ≥50 or 75µm in diameter determined.

### Cell Culture

HEK293T/17 cells were purchased from ATCC (CRL-11268) and were cultured in DMEM (Corning) with 10% FBS. Homozygous MCF7 Y537S *ESR1* cells (generously donated by Dr. Sarat Chandralapaty, MSKCC) were grown in DMEM (Corning) with 5% FBS supplementation. Homozygous T47D Y537S *ESR1* cells (generously donated by Dr. Geoffrey Greene, University of Chicago) were grown in RPMI (Corning) supplemented with 6.5 µg/mL, and 10% FBS. Heterozygous MCF7 Y537S *ESR1* luciferase tagged cells (generously donated by Drs. Geoffrey Greene and Muriel Lainé, University of Chicago) were grown in DMEM (Corning), 5% FBS, and 1% pen-strep and L-glut (Corning). Cells were mycoplasma tested every 15-20 passages and their identities confirmed using STR profiling through ATCC before experiments.

### Scratch Migration Assay

MCF7 Y537S *ESR1* homozygous cells were seeded in a 24 well plate and monitored until 100 percent confluency was achieved. Cells were pretreated with Mitomycin C 2 hours prior to scratch. At this point, a sterile pipette tip was dragged through the center of the well. Immediately after scratch, media was changed and drug was added for a final concentration of 1 µM of T6I-29-1A. Cells were monitored and photos were taken immediately after scratch, and 24 hours after. Distance was measured using imaging software.

### Murine Breast Cancer Models

Murine studies were conducted in compliance with an approved Institutional Animal Care and Use Committee (IACUC) protocols at Loyola University Chicago. Female ovariectomized NOD.Cg-Prkdcscid/J (Jackson Labs) were implanted with 0.30cm silastic capsules containing E2. Mice bilaterally injected with 2 million homozygous MCF7 Y537S *ESR1* cells (generous gift from Dr. Sarat Chandralapaty) in mammary fat pads (pilot mouse study). In subsequent oral pilot study and comparative study with ICI, heterozygous luciferase-labeled MCF7 Y537S *ESR1* cells (generously provided by Dr. Geoffrey Greene) were bilaterally injected into the mammary fat pads at 1, and 1.5 million per mammary fat pad, respectively. In each experiment, cells were individualized and suspended in 100µL of a 1:1 Matrigel (Corning):DMEM (Corning) mixture. Tumor cell growth was monitored via caliper measurements 3x per week. In oral pilot study and comparative study with ICI, tumor cell growth was also monitored with IVIS Spectrum In Vivo Imaging System (Perkin Elmer). To visualize tumor growth, 100µL of 30 mg/mL D-luciferin (PerkinElmer catalog number 122799) suspended in PBS is injected via IP into a 20 gram mouse. After 10 minutes, mice are anestatized with a flow rate of 1-2% and imaged using IVIS Spectrum In Vivo Imaging System (PerkinElmer). Mice were sacrificed when tumor size reached 2000 mm^3^.

### *Ex Vivo* Murine Tissue Imaging

In oral pilot study and comparative ICI mouse studies endpoint, mice were IP injected with 100µL of 30 mg/mL D-luciferin (PerkinElmer catalog number 122799) suspended in PBS. Mice were humanely sacrificed, and relevant tissues including, femurs, lung, liver, uterus, adrenal glands, and brain were imaged rapidly with the IVIS Spectrum In Vivo Imaging System (PerkinElmer). Metastatic burden was calculated by measuring the total flux of each organ (photons per second [p/s]) normalized to average radiance (cm^2^/sr) using Living Image Software (PerkinElmer). Statistical analysis of metastatic burden by organ and overall was performed using one-way ANOVA with relevant post-hoc tests.

### Treatments

In the intraperitoneal (IP) Pilot Murine Xenograft experiment, T6I-29-1A was dissolved in 50 percent DMSO to PBS vehicle at different dosing concentrations (5,10,25,or 100 mg/kg). T6I-29-1A was administered IP 5 times per week (M-F), with tumor caliper measurements performed 3 times per week. In the oral pilot study, T6I-29-1A was dissolved in 0.2 percent tween 80, .5 percent carboxymethylcellulose (CMC) vehicle at different dosing concentrations of 5 or 25 mg/kg. The comparative murine xenograft experiment had three treatment arms, consisting of 10 mice treated with IP vehicle (50% DMSO in PBS) and subcutaneous (SC) vehicle (5% DMSO, 95% peanut oil). 10 mice receiveing T6I-29-1A were treated at 25 mg/kg IP 5 times per week dissolved in IP vehicle, and mice in fulvestrant (ICI) arm received clinically relevant 25 mg/kg dose SC once weekly, as reported as a clinically relevant dose previously [49].

### Histology

Relevant tissues were harvested immediately post-euthanasia and fixed in 10% formalin (Fisher). After 24 hours in formalin, tissues were washed and moved to 70% ethanol in PBS for long term storage and femurs and long bones were decalcified at 4 degrees celcius rocking for 5 days. Hematoxylin and Eosin (H&E) staining was performed by the core at Loyola University Medical Center (LUMC) by Lourdcymole Pazhampally for the IP pilot mouse study and uterine wet weight studies. All other H&E staining was performed using Eprendia Hematoxylin (catalog number 7211) and eosin (catalog number 7111). Slides were analyzed for metastatic lesion analysis by Dr. Khin Su Mon (LUMC pathology) in the IP pilot study and Dr. Marteen Bosland (UIC pathology) for subsequent studies.

### Uterine Wet Weight Study

Murine studies were conducted in compliance with an approved Institutional Animal Care and Use Committee (IACUC) protocols at Loyola University Chicago. Adult female ovariectomized BALB/c mice (Jackson labs) were assigned randomly to 8 groups. Mice groups were treated once daily with one of the following: vehicle (0.2 percent tween 80, .5 percent CMC), E2 (0.1mL of 0.1 µg/mL E2 in 95% cottonseed oil and 5% ethanol), tamoxifen (50 mg/kg tamoxifen in vehicle), tamoxifen +E2, ICI (25 mg/kg in 95% cottonseed oil and 5% ethanol), ICI+E2, T6I-29-1A (50 mg/kg orally dosed), or T6I-29-1A +E2. After three days of consecutive treatment, animals were humanely euthanized and uteri were weighed and embedded and fixed in cassettes [30].

### NanoBiT ERα-SRC3 assay

HEK293T/17 cells were grown in a white walled, 96-well clear bottom plate, seeded at 9k/well. When cells achieved 50-70% confluence, they were transfected with DNA plasmids containing C-terminus tagged smBiT ERα, smBiT ERα Y537S and N-terminus tagged lgBiT SRC3 were generously donated by Dr. Donald P. McDonnell. smBiT was cotransfected with lgBiT SRC-3 at 0.1 µg/ plasmid per well with 3:1 µL:µg Turbofectin 8.0 (Origene, TF81001) with 9µL Opti-MEM/well (Gibco/ Thermo Fisher, catalog number 31-985-070) in full media. After 24 hours, media was replaced with serum-starved media for 72 hours. Cells were then treated in the presence of 1 nM E2 with different compounds at various concentrations, with 1% vehicle (DMSO) and 1 nM E2 controls on each plate. Cells were treated with Nano-Glo substrate (Promega, catalog number N2012) at a 1:20 dilution in buffer and read immediately for luminescent signal using BioTek Cytation 5. The data shown are three biological replicates, with nine total replicates per concentration.

### Cell Proliferation

MCF7 Y537S *ESR1* and T47D Y537S *ESR1* cells were seeded at low confluencies (750 cells per well and 1000 cells per well, respectively) in serum-starved media on a 96 well plate. Cells were in serum starved media for 72 hours prior to treatment. After 72 hours, cells were treated with vehicle (1% DMSO), 1 nM E2, or 1 µM SERM/ SERD +1 nM E2. Cells were grown in a BioSpa attached to a BioTek Cytation 4 and were automatically counted with the BioTek software every 12 hours for 5-10 days, or until the E2-only wells reached confluency. Media and drug treatment were replaced every 3-4 days. Each graph represents three replicates with three separate repeats.

### RNA sequencing

Homozygous MCF7 Y537S *ESR1* breast cancer cells were grown in 6 well dishes in serum-starved media for 48 hours. Upon reaching 50 percent confluency, cells were treated with vehicle (1% DMSO), 1 nM E2, or 1 nM E2 + 1 µM ICI, Laso, Rad1901, or T6I-29-1A for 24 hours in triplicate. After 24 hours, RNA was isolated using Quigen RNeasy Kit and sent to Novogene for sequencing and bioinformatics analysis.

### ELISA assays

ELISA plates from Thermo Fisher (catalog number 12-565-135) were filled with 100uL of standard per well or plasma sample dilution. Each sample was ran in duplicate and each standard curve was run in duplicate on each plate. Recombinant DKK1 was purchased from Gibco through Fisher (catalog number PHC9214). Recombinant DKK1 was reconstitutied as per manufacturer instructions and further diluted in 2 mg/mL BSA in PBS. Standard curve dilutions ranged from 30000 pg/mL to 122.9 pg/mL. 1 to 100 dilution of plasma samples was used (as it was determined to be in linear range) to interpolate DKK1 values. Plasma samples were diluted in 2 mg/mL BSA in PBS for dilutions. DKK1 was detected using DKK1 monoclonal rabbit antibody (Invitrogen, 1D12), and detected with secondary antibody conjugated to HRP (Fisher, PI31460) followed by incubation with TMB substrate (Fisher, ENN301). Reaction was quenched with diluted sulfuric acid. Absorbance was read on BioTek Cytation 5 plate reader at 450nm.

### Healthy and ER+ Breast Cancer Patient Plasma Samples

Healthy women plasma samples were obtained through the Komen Tissue Bank (KTB) at Indiana University. ER+ Breast Cancer Patient Plasma and chart review was obtained through the Indiana University Simon Comprehensive Cancer Center (IUSCCC).

### Quantitative PCR (qPCR)

MCF7 Y537S *ESR1* or T47D Y537S *ESR1* breast cancer cells were seeded in 6-well plates in serum starved media for 72 hours. After 72 hours, cells were confirmed to have reached 50-70 percent confluence and were treated with 1 nM E2 and 1 µM of each SERM or SERD indicated. After 24 hours, RNA was harvested using the Qiagen RNeasy Kit. cDNA was made using the M-MLV reverse transcriptase (Invitrogen, catalog number 28025013). qPCR was performed using Power up SYBR Green Master Mix (Thermo Fisher, catalog number A25741).

### Primers

*GREB1* F: 5′-CTGCCCCAGAATGGTTTTTA-3′

*GREB1* R: 5′-GGACTGCAGAGTCCAGAAGC-3′

*PGR* F: 5′-AGCCAGAGCCCACAATACAG-3′

*PGR* R: 5′-GACCTTACAGCTCCCACAGG-3′

*CA12* F: 5′-GACCTTTATCCTGACGCCAGCA-3′

*CA12* R: 5′-CATAGGACGGATTGAAGGAGCC-3′

*cMyc* F: 5′-TTCGGGTAGTGGAAAACCAG-3′

*cMyc* R: 5′-CAGCAGCTCGAATTTCTTCC-3′

## Acknowledgments

This work was funded by the National Cancer Institute, National Institutes of Health R37CA279341, Susan G. Komen CCR19608597, and a seed grant from the Cardinal Bernardin Cancer Center to S.W.F. This research used resources from the 17-ID-1 AMX beamline of the National Synchotron Light Souce II, a U.S. Department of Energy (DOE) Office of Science by Brookhaven National Laboratory under Contract No. DE-SC0012704. The Center for BioMolecular Structure (CBMS) is primarily supported by the National Institutes of Health, National Institute of General Medical Sciences (NIGMS) through a Center Core P30 Grant (P30GM133893), and by the DOE Office of Biological and Environmental Research (KP1605010).

## Notes

### Competing Interest Statement

Sean Fanning has patent on T6I-29. PCT/US2022/016813 Estrogen Receptor Alpha Antagonists and Uses Thereof.

