## Supplemental Files for "Targeting Unique Ligand Binding Domain Structural Features Downregulates DKK1 in Y537S *ESR1* Mutant Breast Cancer Cells"

### SUPPLEMENTAL FIGURES

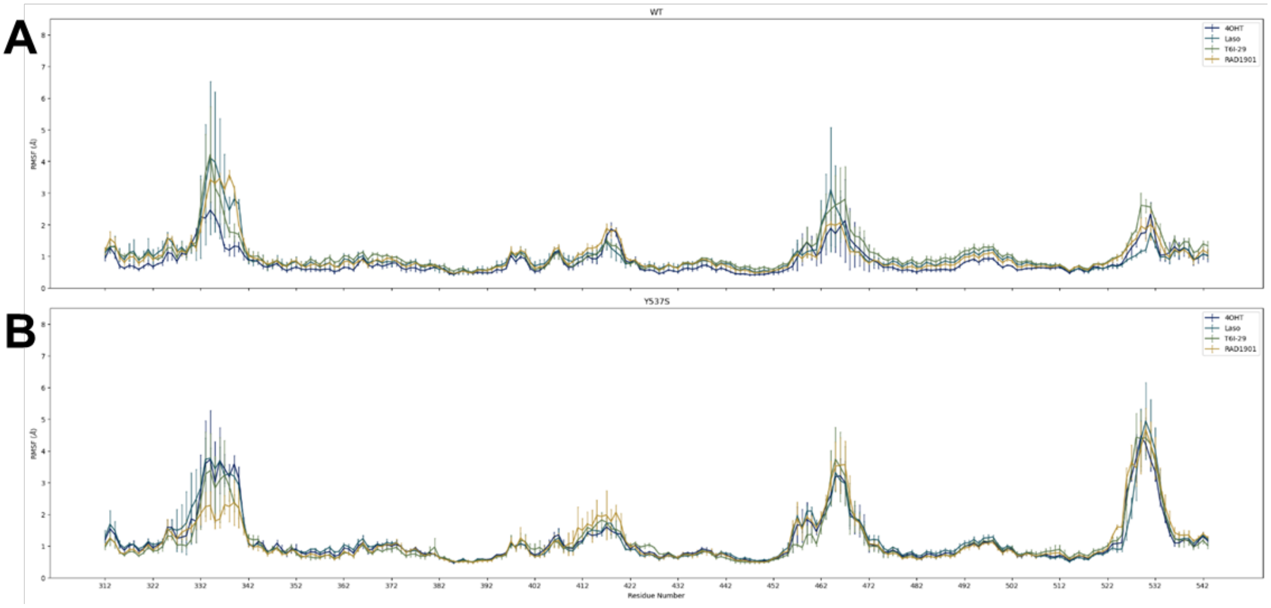


**Supplemental Figure 1:** Average RMSF per residue over triplicate simulations of each ligand bound to **A)** WT or **B)** Y537S ERα LBD.

| PK parameters | Unit | Mouse 1 | Mouse 2 | Mouse 3 | Mean | SD | CV |
|---|---|---|---|---|---|---|---|
| $T_{1/2}$ | h | 3.16 | 5.07 | 3.84 | 4.02 | 0.96 | 24.0 |
| $T_{max}$ | h | 1.00 | 1.00 | 1.00 | 1.00 | 0.00 | 0.000 |
| $C_{max}$ | ng/mL | 771 | 674 | 811 | 752 | 70 | 9.37 |
| $AUC_{last}$ | h*ng/mL | 2478 | 2757 | 3431 | 2888 | 490 | 17.0 |
| $AUC_{Inf}$ | h*ng/mL | 2485 | 2831 | 3477 | 2931 | 503 | 17.2 |
| $AUC_{_{\%Extrap}}$_obs | % | 0.303 | 2.61 | 1.33 | 1.41 | 1.15 | 81.8 |
| $MRT_{Inf}$_obs | h | 2.99 | 4.48 | 4.17 | 3.88 | 0.78 | 20.2 |
| $AUC_{last}$/D | h*mg/mL | 99.1 | 110 | 137 | 116 | 20 | 17.0 |
| F | % | NA | NA | NA | NA | NA | NA |

**Supplemental Table 1:** PO DMPK of T6I-29.

| PK parameters | Unit | Mouse 4 | Mouse 5 | Mouse 6 | Mean | SD | CV |
|---|---|---|---|---|---|---|---|
| $T_{1/2}$ | h | 3.58 | 3.57 | 3.70 | 3.61 | 0.07 | 1.97 |
| $T_{max}$ | h | 0.500 | 0.500 | 0.500 | 0.500 | 0.000 | 0.000 |
| $C_{max}$ | ng/mL | 4420 | 4540 | 6200 | 5053 | 995 | 19.7 |
| $AUC_{last}$ | h*ng/mL | 7355 | 8195 | 9413 | 8321 | 1035 | 12.4 |
| $AUC_{Inf}$ | h*ng/mL | 7381 | 8222 | 9446 | 8350 | 1038 | 12.4 |
| AUC_%Extrap_obs | % | 0.349 | 0.333 | 0.353 | 0.345 | 0.010 | 2.99 |
| $MRT_{Inf}$_obs | h | 2.11 | 2.28 | 2.21 | 2.20 | 0.08 | 3.70 |
| $AUC_{last}$/D | h*mg/mL | 294 | 328 | 377 | 333 | 41 | 12.4 |
| F | % | NA | NA | NA | NA | NA | NA |

**Supplemental Table 2:** IP DMPK of T6I-29.

| | Y537S-T6I-4-1 | Y537S-T6I-14-1 | Y537S-T6I-29-1A |
|---|---|---|---|
| PDB ID | 9BU1 | 9BQE | 9BPX |
| Data Collection | | | |
| Space Group | C2 | C2 | C2 |
| Cell Dimensions | | | |
| a, b, c (Å) | 102.75, 57.98, 87.62 | 103.18, 56.94, 87.72 | 101.73, 58.29, 87.01 |
| α, β, γ (°) | 90.00, 103.95, 90.00 | 90.00, 104.53, 90.00 | 90.00, 103.52, 90.00 |
| Resolution (Å) | 1.75 | 1.98 | 2.20 |
| $CC^{1/2}$ | 0.998 (0.642) | 0.998 (0.640) | 0.999 (0.655) |
| I/σI | 2.08 (at 1.75 Å) | 1.24 (at 1.98 Å) | 2.96 (at 2.20 Å) |
| Completeness | 99.2 | 98.6 | 98.1 |
| Redundancy | 3.7 | 3.7 | 3.7 |
| Refinement | | | |

| | | | |
|---|---|---|---|
| Resolution Range (Å) | 20 – 1.75 Å | 20 – 1.98 Å | 20 – 2.20 Å |
| Number of Reflections | 50,328 | 37,473 | 24,826 |
| $R_{work}/R_{free}$ | 19.4/22.6 | 21.7/26.7 | 18.6/22.1 |
| No. Atoms | 3,883 | 3,577 | 3,917 |
| Water Molecules | 423 | 121 | 186 |
| Ligand Molecules | 2 | 2 | 2 |
| R.m.s. deviations | | | |
| Bond Lengths (Å) | 0.007 | 0.003 | 0.004 |
| Bond Angles ($^{o}$) | 1.39 | 0.597 | 0.600 |
| Ramachandran | | | |
| Favored (%) | 98.77 | 98.56 | 97.19 |
| Allowed (%) | 1.23 | 1.44 | 2.81 |
| Outliers (%) | 0 | 0 | 0 |

**Supplemental Table 3:** X-ray data collection and refinement statistics.

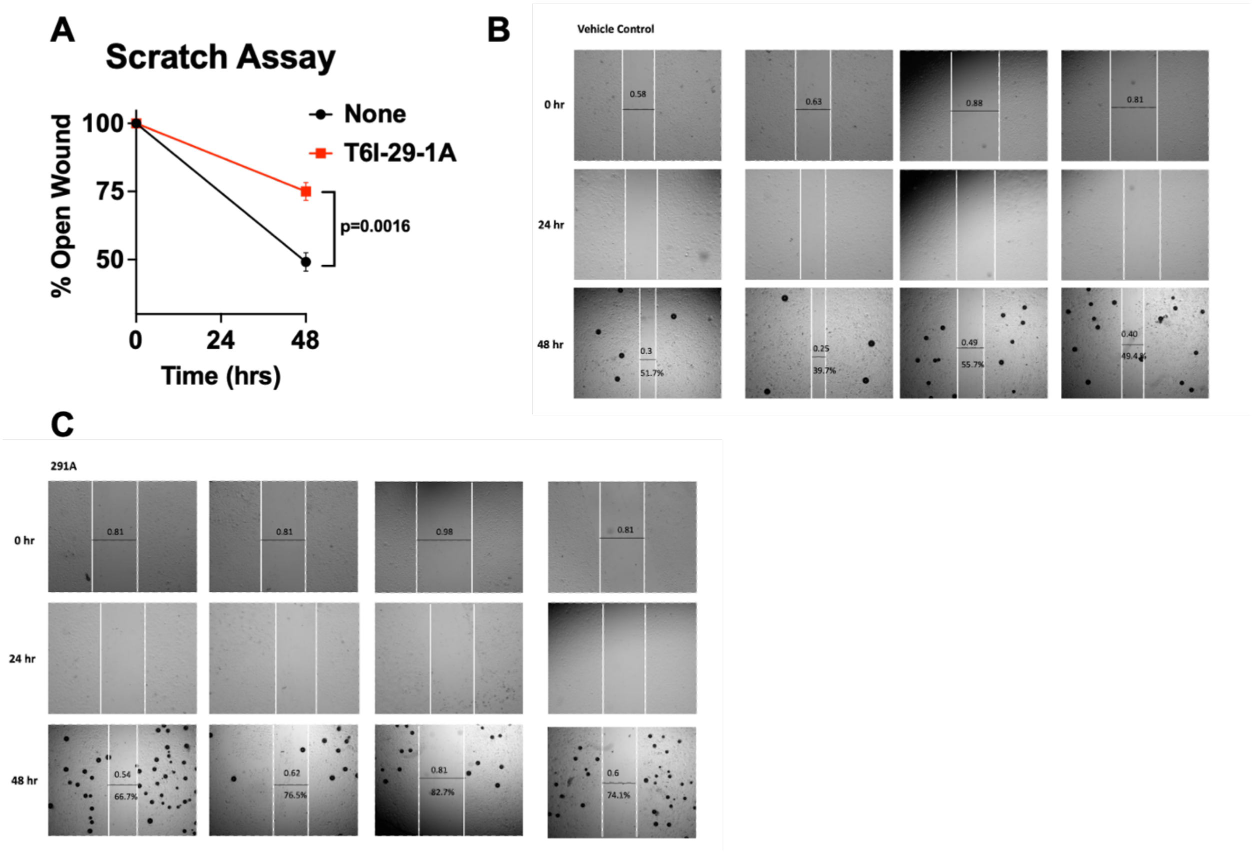


**Supplemental Figure 2:** T6I-29-1A significantly blunts migratory capacity of MCF7 Y537S *ESR1* cells. A) Quantified open wound percentage using imaging software. Significance is determined using unpaired T test. B-C) Representative images of cells treated with vehicle (B) and T6I-29-1A (C).

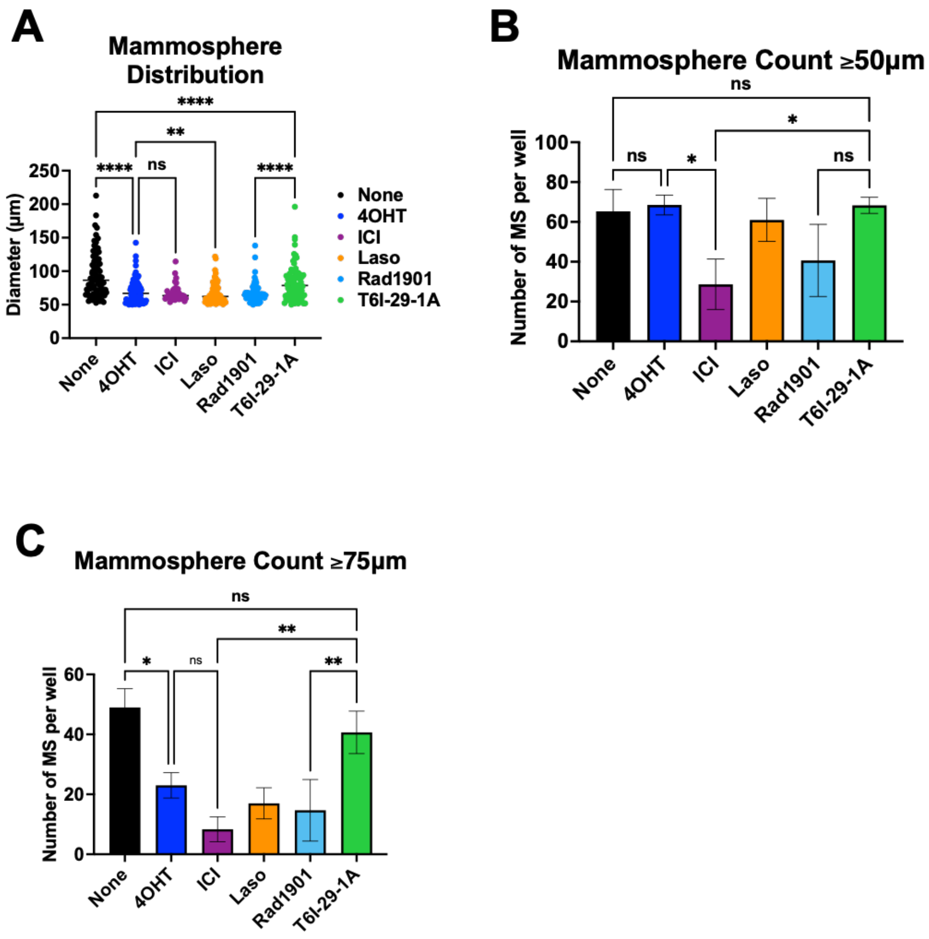


**Supplemental Figure 3**: T6I-29-1A decreases mammosphere size, but does not change total count. A) Mammosphere size quantified using imaging software, significance is determined by ANOVA with Tukey post-hoc test, *p < 0.05, **p < 0.005, ***p < 0.0005, and ****p < 0.00005. B) Number of mammospheres that are greater or equal in size of 50 µm, same statistical tests used as A. C) Number of mammospheres that are greater or equal in size to 75 µm. Same statistical tests used as A and B.

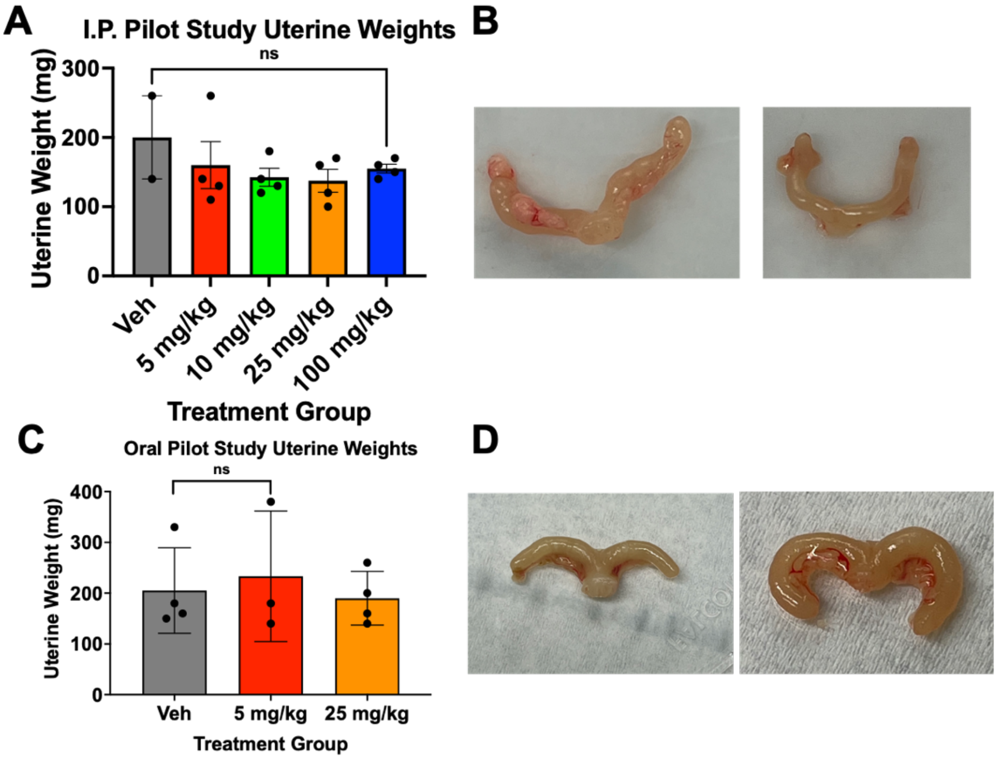


**Supplemental Figure 4:** T6I-29-1A does not significantly stimulate uterines in pilot studies. *Ex vivo* uterine weight taken by digital scale in A) pilot IP study and B) oral pilot study with representative uterine photos respectively in B, D. Statistical analysis performed by unpaired t test.

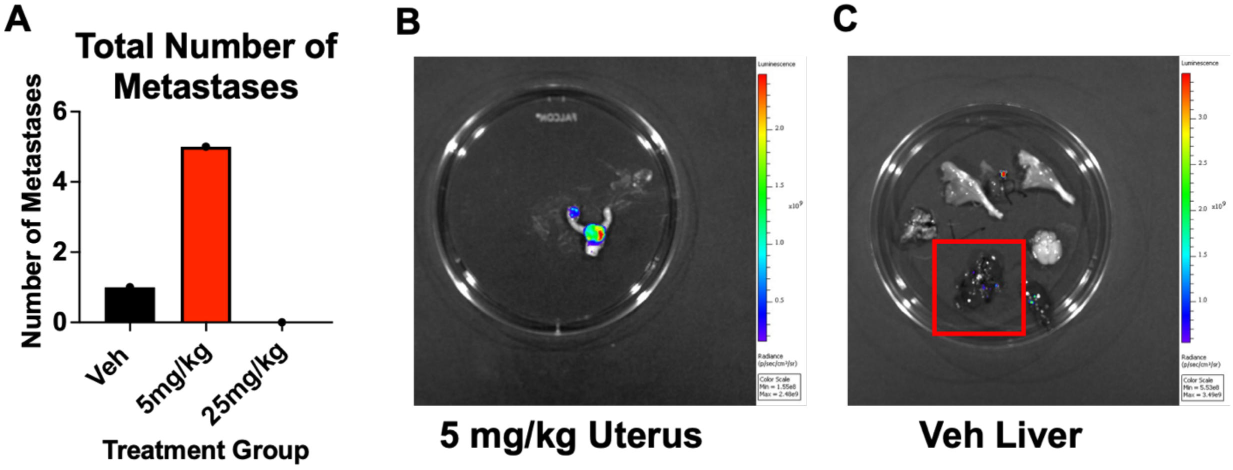


**Supplemental Figure 5:** Metastatic burden in oral pilot study measured from H&E slides, confirmed with IVIS *ex vivo* imaging. A) Total number of metastases measured by pathologist Dr. Marteen Bosland across all treatment groups. B) Uterus metastatsis confirmed by IVIS in 5 mg/kg T6I-29-1A treatment group. C) Red rectangle shows small metastatses in liver that were noted in H&E slide analysis; confirmed by IVIS.

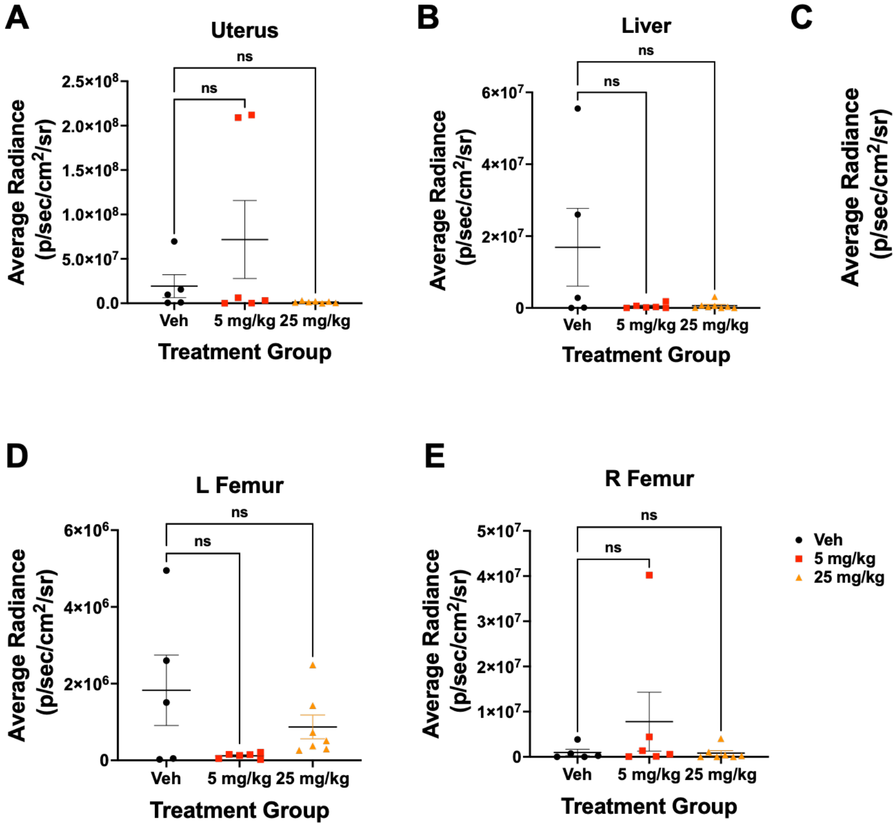

**Supplemental Figure 6:** *Ex vivo* imaging shows 25 mg/kg dose attenuates metastates in brain, and has differential effects on other organs metastatic colonization. IVIS Spectrum imaging of front and back of A) Uterus, B) Liver, C) Brain, D) L Femur and E) R Femur show trends in attenuation of metastatses compared to vehicle, with a significant decrease in brain metastases, in the 25 mg/kg oral dose (C), **p < 0.005. Error bars represent SEM.

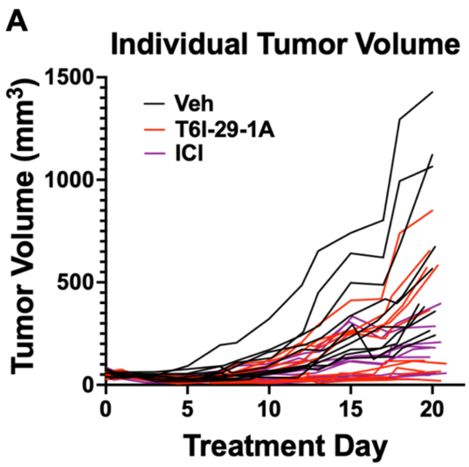


**Supplemental Figure 7:** Individual mouse tumor growth shows T6I-29-1A treated mice situated between growth of vehicle and ICI treated mice. A) Tumor volume as measured by digital caliper three times per week. Each line on graph represents a separate mouse tumor growth.

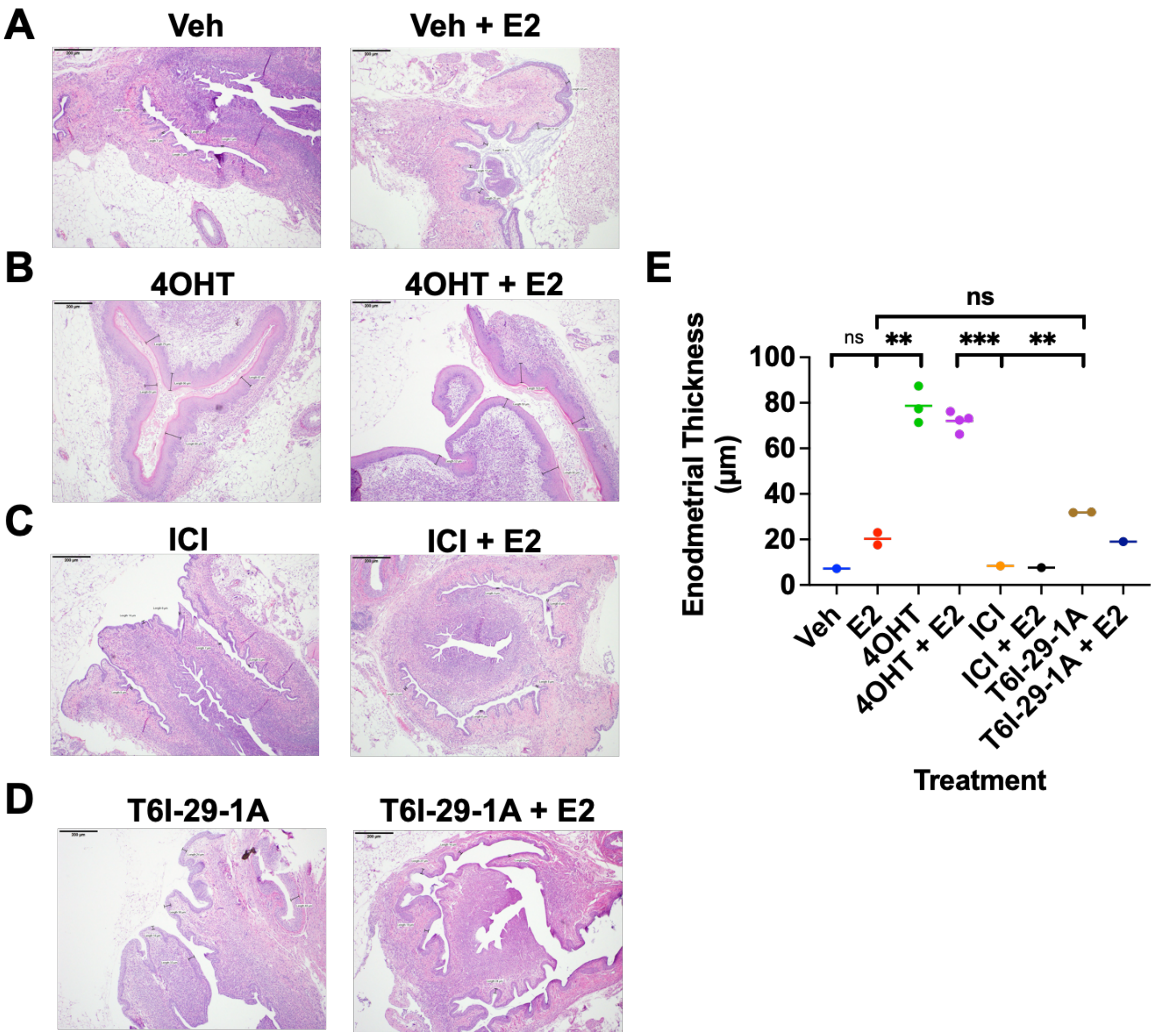


**Supplemental Figure 8:** Uterine wet weight study suggests decreased endometrial thickness with ICI treatment. A-D) Representative images of Vehicle treated (A), Tamoxifen treated (B), ICI treated (C), and T6I-29-1A treated (D) uteri, visualized by H&E staining. Representative measurements of endometrial thickness are shown, with an average of at least three measurements taken for analysis. Measurements taken with imaging software ImageJ. E) Enodmetrial thickness from an average of at least three separate measurements per uterus. Statistical analysis conducted using ANOVA with relevant post-hoc test.

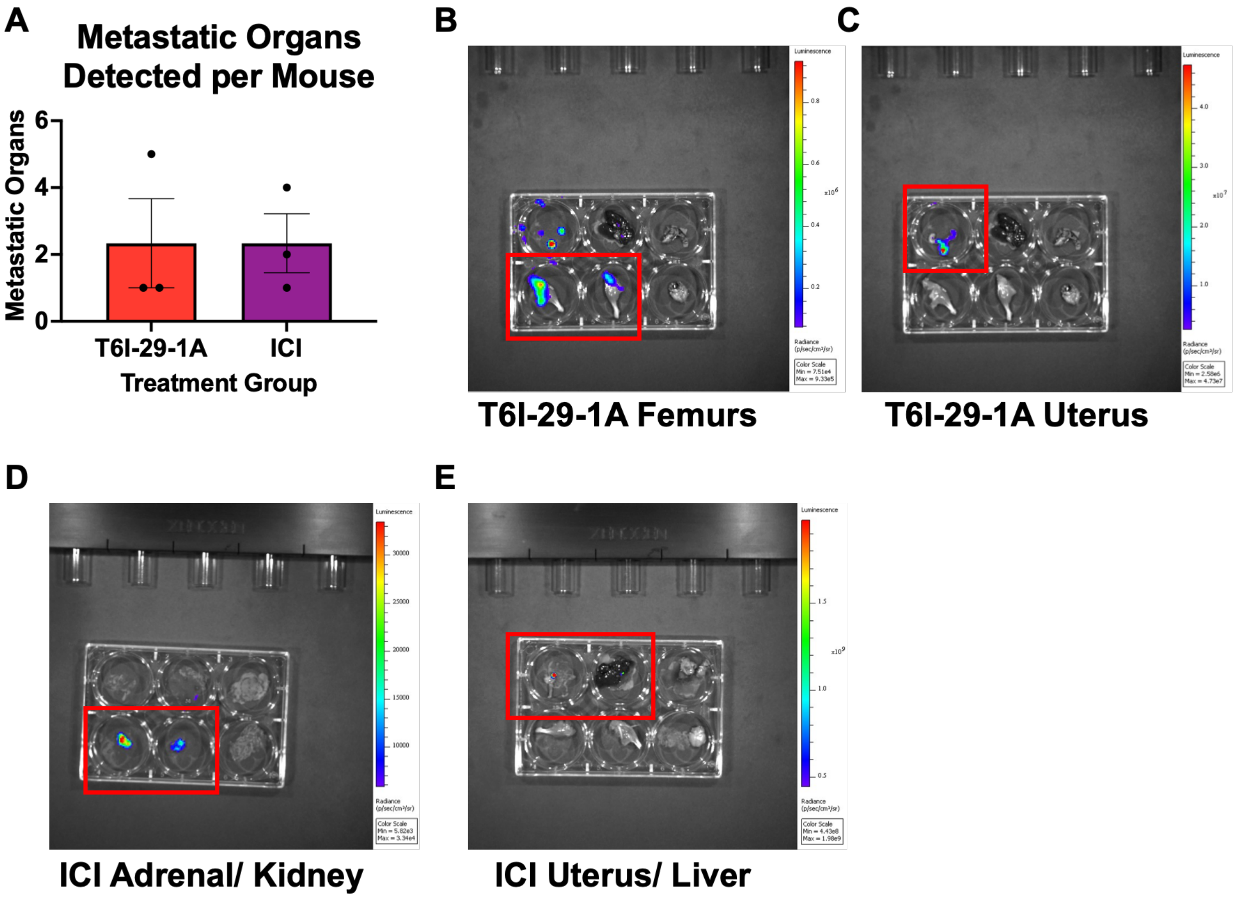


**Supplemental Figure 9:** Comparative study metastatic lesions measured by H&E staining, confirmed by IVIS *ex vivo* imaging. A) Organs that had one or more metastatic lesion are counted on a per mouse basis. No lesions were found in vehicle mice via H&E pathological analysis conducted by Dr. Marteen Bosland. Red box denotes B) Femur metastases, C) Uterus, D) Adrenal/ Kidney and E) Uterus/ Liver confirmed by IVIS that were identified by H&E staining in T6I-29-1A (B,C) and ICI (D,E) treatment groups.

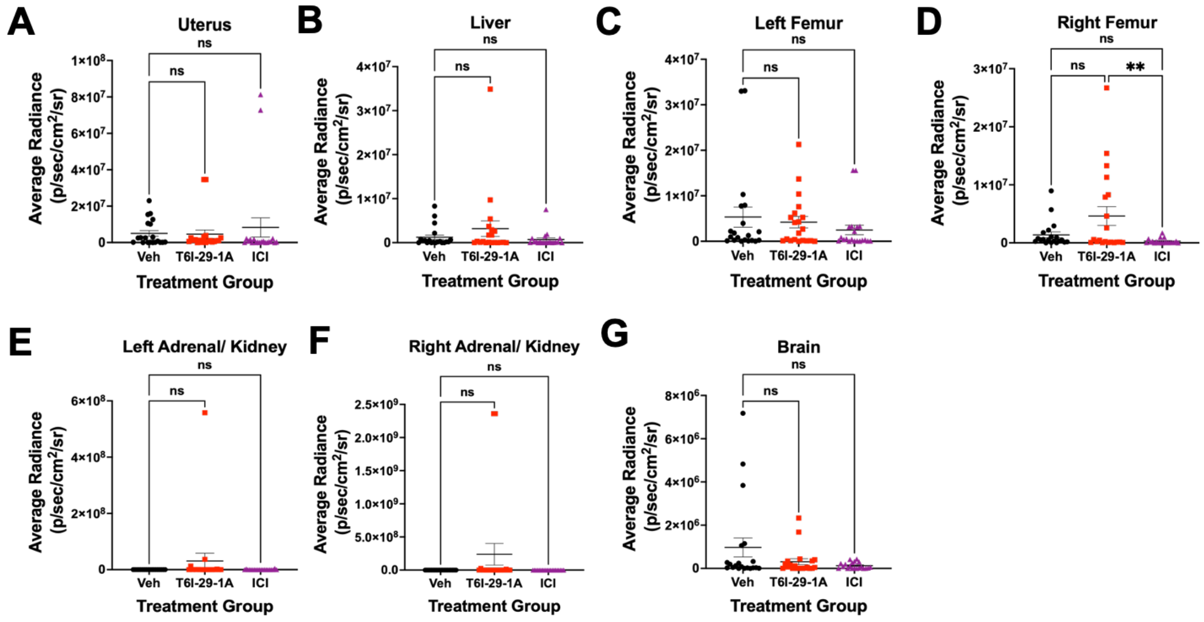


**Supplemental Figure 10:** *Ex Vivo* Imaging shows trends in differences in metastatic colonization based on treatment group. Common metastatic sites for MCF7 Y537S *ESR1* breast cancer cells are analyzed *ex vivo* for each mouse and results compiled for A) Uterus, B) Liver, C-D) Femurs, E-F) Left and Right Adrenal/ Kidneys, and G) Brain. Statistical analysis performed using ANOVA with Tukey post hoc test.

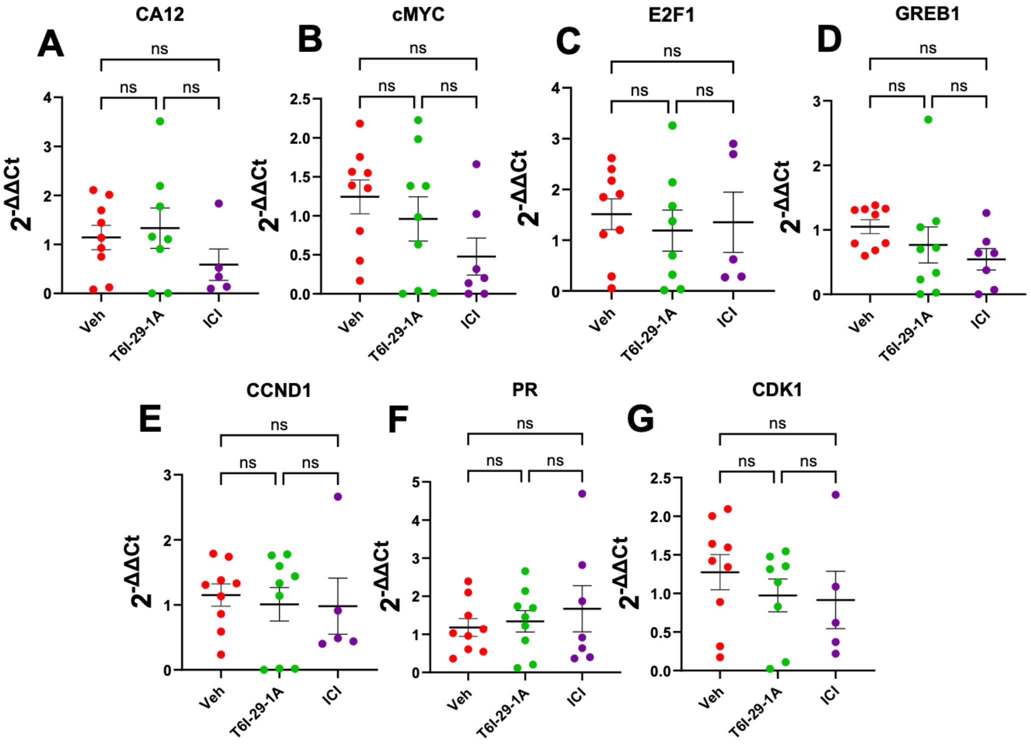

**Supplemental Figure 11:** qPCR on MCF7 Y537S *ESR1* tumors shows trends in ER target gene downregulation. Common ER target genes were analyzed using qPCR on xenograft tumors. A-E, G) *CA12, cMYC, E2F1, GREB1, CCND1,* and *CDK1* ER target genes show general trends in downregulation from vehicle, to T6I-29-1A, to ICI treated mice. F) *PR* does not appear regulated by T6I-29-1A or ICI treatment *in vivo.* Statistical analyses performed using ANOVA with Tukey post-hoc test.

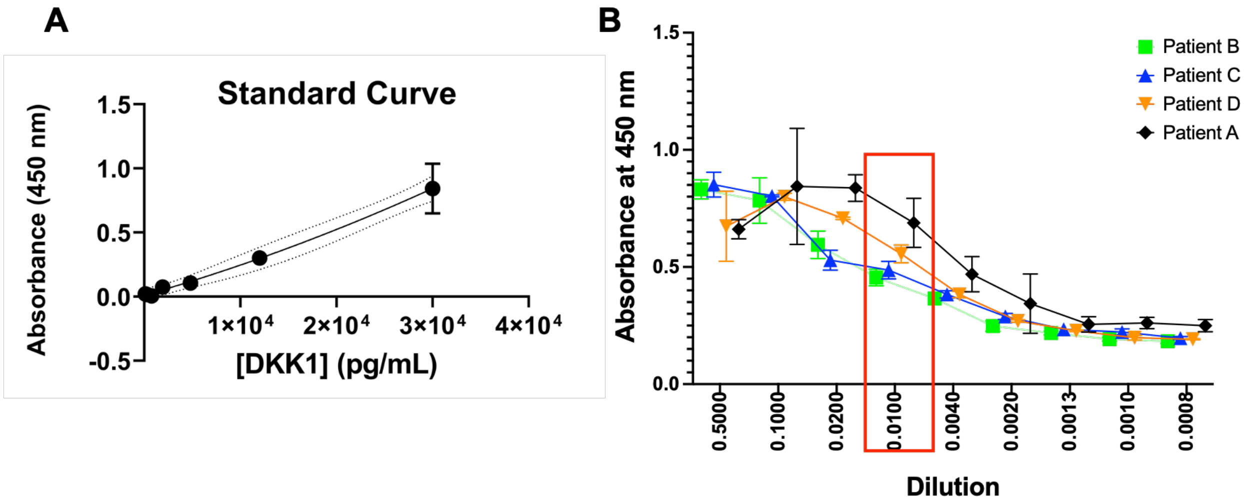


**Supplemental Figure 12:** DKK1 standard curve and patient dilution curve employed to find accurate linear range. A) Recombinant DKK1 at different concentrations was ran on every ELISA plate to determine standard curve. Standard curves were fit with a polynomial curve, with R squared values over .9. B) Example dilution range for patient plasma ran in duplicate on ELISA plate. 1 to 100 dilution was found to be optimal for patients.
